# A comprehensive atlas of white matter tracts in the chimpanzee

**DOI:** 10.1101/2020.01.24.918516

**Authors:** Katherine L. Bryant, Longchuan Li, Nicole Eichert, Rogier B. Mars

## Abstract

Chimpanzees (*Pan troglodytes*) are, along with bonobos, humans’ closest living relatives. The advent of diffusion MRI tractography in recent years has allowed a resurgence of comparative neuroanatomical studies in humans and other primate species. Here we offer, in comparative perspective, the first chimpanzee white matter atlas, constructed from *in vivo* chimpanzee diffusion-weighted scans. Comparative white matter atlases provide a useful tool for identifying neuroanatomical differences and similarities between humans and other primate species. Until now, comprehensive fascicular atlases have been created for humans (*Homo sapiens*), rhesus macaques (*Macaca mulatta)*, and several other nonhuman primate species, but never in a nonhuman ape. Information on chimpanzee neuroanatomy is essential for understanding the anatomical specializations of white matter organization that are unique to the human lineage.

## Introduction

Knowledge about chimpanzee brains is critical to understanding the neural basis of the unique behavioral adaptations of both humans and chimpanzees. Until recently, there were few options for examining the organization of chimpanzee brains, as methods used on other primate species, such as invasive tract-tracing, are not possible for ethical reasons. However, the advent of non-invasive structural neuroimaging methods has opened up a new era of comparative n
euroanatomy. Using such tools, white matter atlases have been produced for the human [1,2], macaque monkey [3], and squirrel monkey [4]. Although a few investigations into the anatomy of selected fasciculi in chimpanzees have been undertaken [5–7], thus far no study has reconstructed all of the major fasciculi. Here, we take up this challenge and provide a first comprehensive white matter atlas for the chimpanzee, who, along with the bonobo, is our closest living animal relative.

We identify the major white matter fibers by means of standardized anatomical landmarks that can be directly compared to those of previous studies in the human and macaque monkey brain [8]. These landmarks are used to create ‘recipes’, or protocols, for each tract in a standardized brain template: a group of seed, target, and exclusion region-of-interest masks (ROIs) for each fasciculus, which, when used together for tractography, reconstruct the desired tract [8,9]. The goal of these recipes is to make them general enough to identify a given tract in every individual, but, through the specific combination of inclusion and exclusion masks, also selective to the tracts of interest. These recipes can be easily transformed to the space of individual scans in different datasets. We use these recipes to reconstruct the major white matter tracts in 29 *in vivo* diffusion-weighted MRI datasets using probabilistic tractography [10]. The nature of these recipes also means that future modifications can easily be incorporated into the atlas, and interested researchers can test out competing recipes to compare claims between rival definitions of any given tract. A specialized tool for implementing these recipes, compatible with our data, has recently been released [11].

We discuss some interesting similarities and differences in the organization of major fasciculi, and their patterns of cortical terminations. Further, we offer resources for neuroanatomists interested in the evolution of the human and chimpanzee brain: an atlas of 24 major white matter tracts in chimpanzees, with directly comparable tracts in humans and macaques, surface projection data from these tracts, and tractography “recipes” in standard space for reproducing these tracts. These resources are made available to the scientific community in online repositories (www.neuroecologylab.org).

## Results

### Overview

We created tractography recipes for 24 tracts in the chimpanzee brain as well as corresponding recipes for the human and macaque (Figs. 1-3). These recipes, described in detail in the Methods section, were used to reconstruct 24 major tracts in the chimpanzee, and, for comparison, human and macaque (Figs. 4-11).

**Figure 1.**
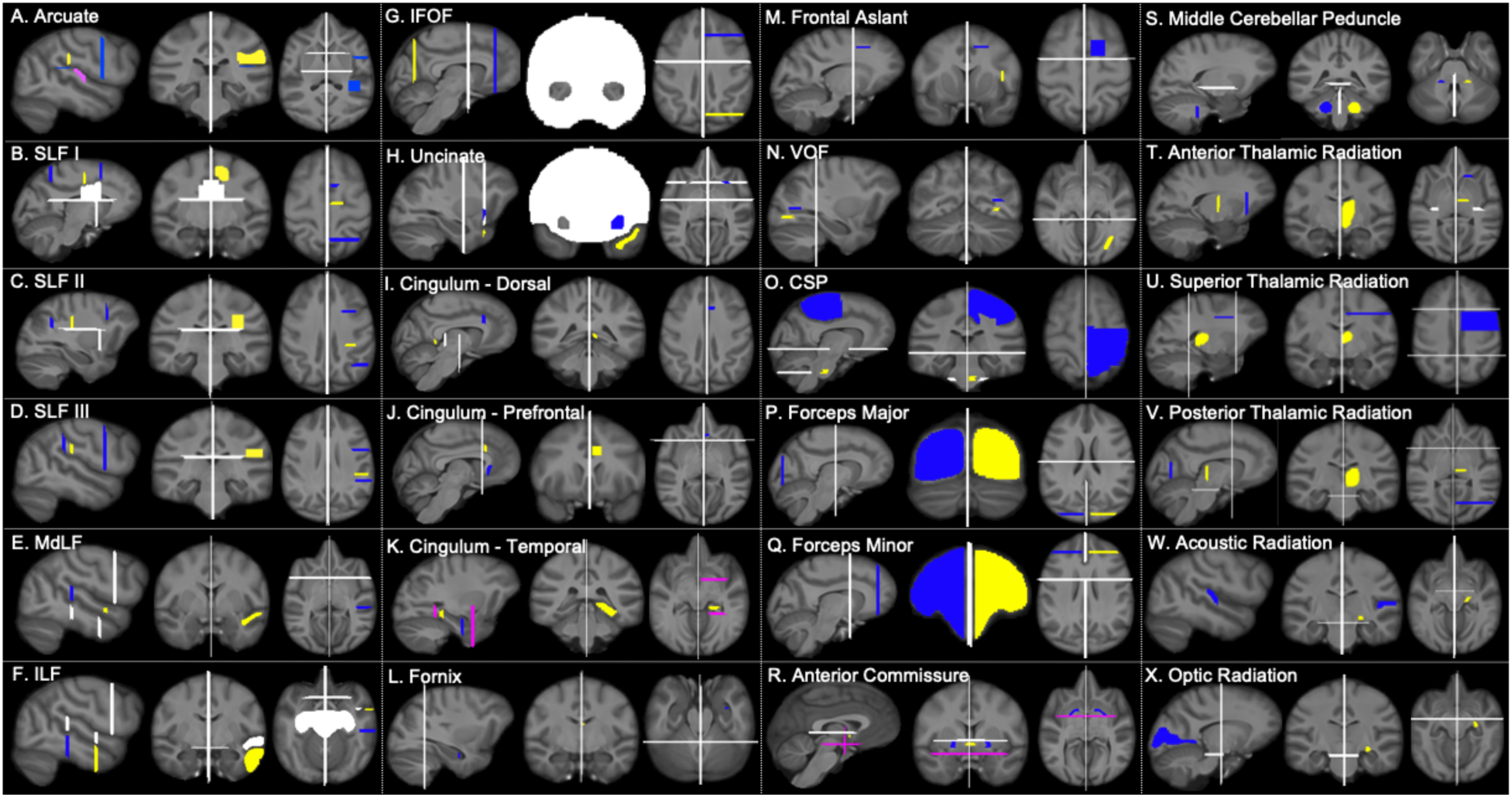
Chimpanzee tractography recipes. Seed ROIs (yellow), target ROIs (dark blue), exclusion masks (white), stop masks (fuschia). Left hemisphere protocols are displayed.

**Figure 2.**
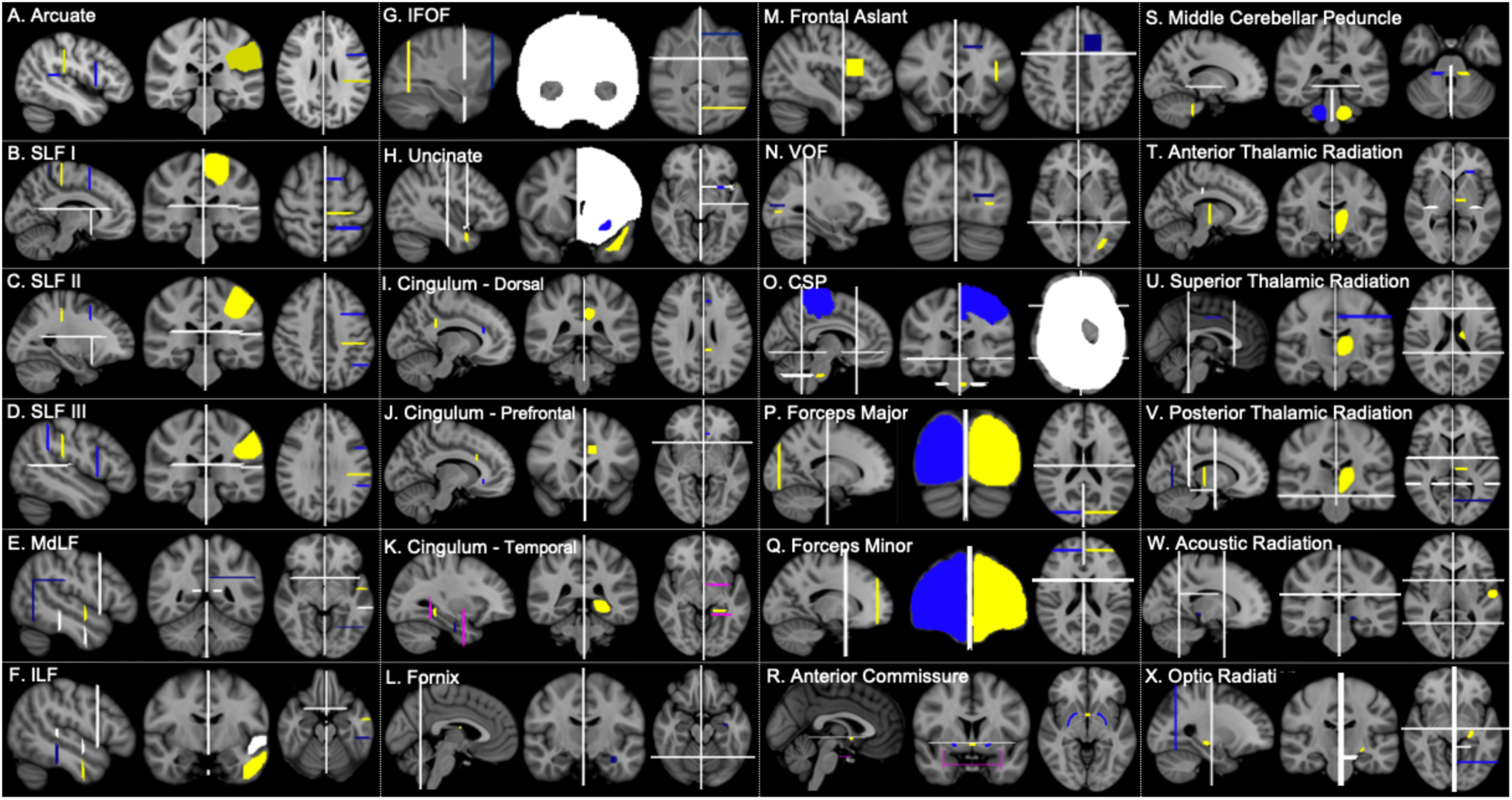
Human tractography recipes. Seed ROIs (yellow), target ROIs (dark blue), exclusion masks (white), stop masks (fuschia). Left hemisphere protocols are displayed.

**Figure 3.**
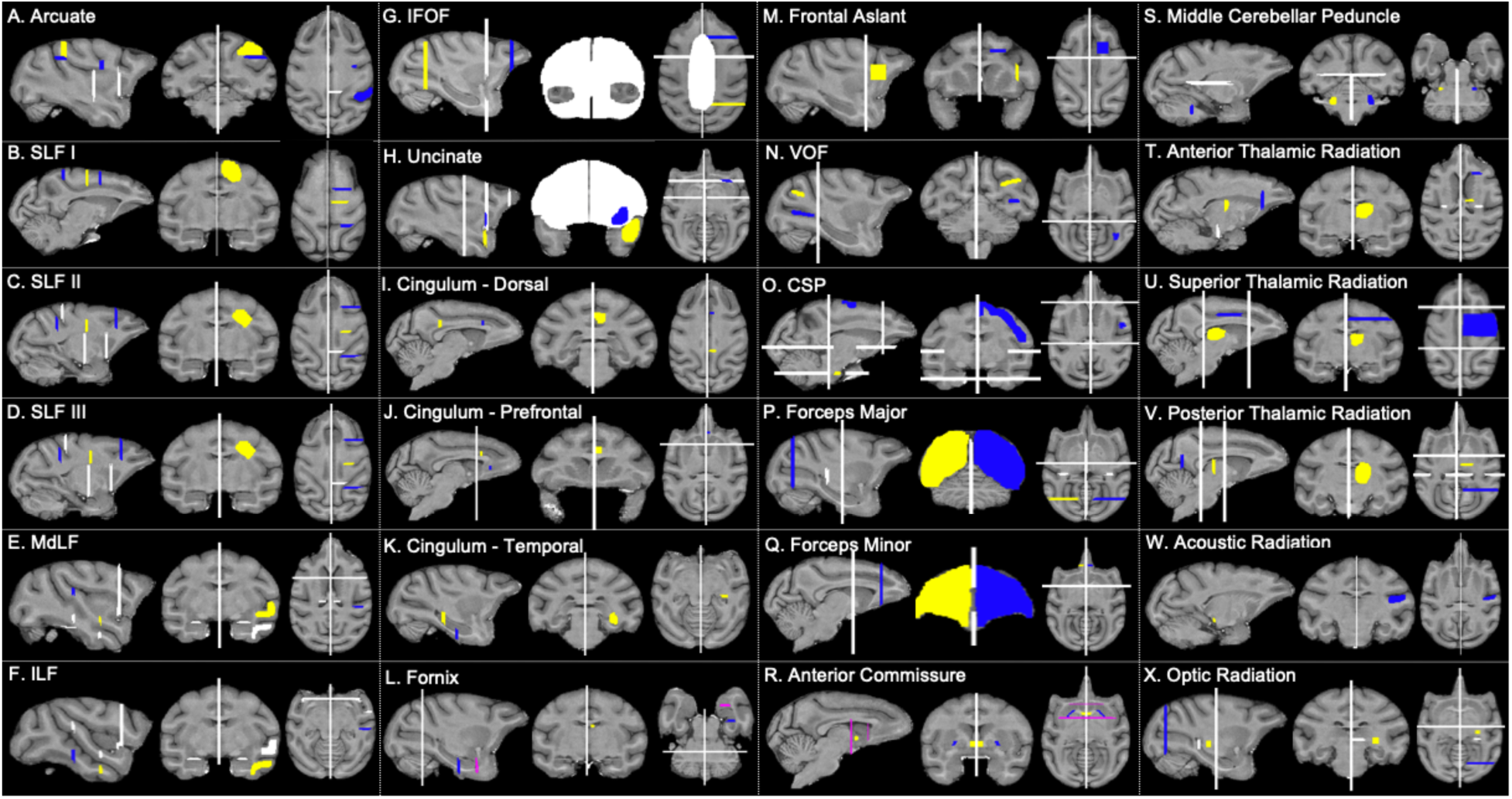
Macaque tractography recipes. Seed ROIs (yellow), target ROIs (dark blue), exclusion masks (white), stop masks (fuschia). Left hemisphere protocols are displayed.

**Figure 4.**
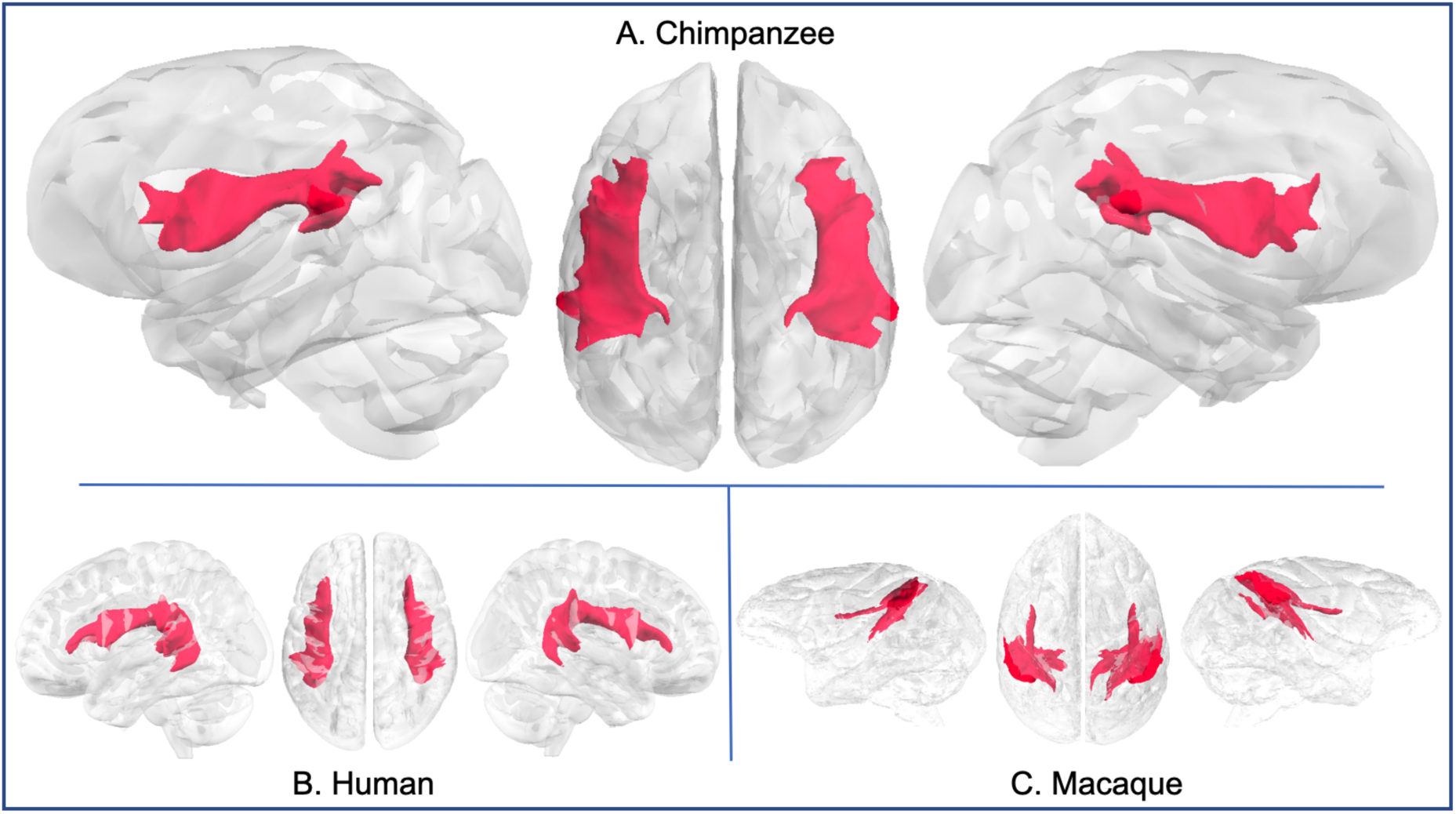
Arcuate fascicle in chimpanzee (a), human (b), and rhesus macaque (c).

The focus of our approach is to identify the bodies of tracts, which can be reliably identified using tractography. Importantly, tractography allows one to reconstruct large white matter fiber bundles, but does not necessarily respect synaptic boundaries. Thus, results obtained using this approach might best be compared to those obtained using blunt dissections, rather than tracer data. This approach of reconstructing tract bodies have in the past proven robust, e.g. [8,12,13], and does not suffer from the disadvantages commonly associated with tractography approaches that aim to mimic tracer data [14,15]. We discuss the course of the tract bodies below and point out important differences between the chimpanzee and the other two species.

For illustration purposes, we also provide surface projection maps (supplemental figs. S1-S5). Due to the nature of tractography, following the course of a fiber bundle as it enters the grey matter presents challenges [14]. We therefore use an alternative approach, of multiplying the tract body with a surface-to-white matter tractogram, rather than following the tract body into the grey matter. This approach has proven to yield replicable and meaningful results [8,11], but we emphasize that the projection maps should not be interpreted as neurocartographic representations of synaptic connections with cortex but rather as depictions of broader trends in fasciculo-cortical connectivity.

A note about terminology: For observed synapomorphies (shared derived characters) between humans and chimpanzees, we have chosen to take a node-based approach [16] by referring to our ingroup with the term “hominin”, referring to the clade including humans, chimpanzees, and bonobos.

### Dorsal longitudinal tracts

The dorsal longitudinal fibers connecting the frontal lobe with the parietal and posterior temporal cortices are formed by the three branches of the superior longitudinal fascicle (SLF) and the arcuate fascicle. We here follow the convention of Schmahmann and Pandya [17] of considering these as distinct tracts, even though the names have been used interchangeably in the literature. All four tracts have been identified using tractography in the human [13,18,19]. They can also be identified using tractography in the macaque [3,13], but appear weaker than in the human, e.g. [8,20].

The chimpanzee arcuate tractogram reached posterior superior temporal gyri, inferior parietal lobule, and posterior inferior frontal gyrus (IFG; Fig. 4a). Compared with chimpanzees, the human arcuate tractogram reached further inferiorly in the temporal lobe (Fig. 4b), consistent with previous findings [6,20]. In contrast, macaque arcuate demonstrated weaker frontal projections than in the chimpanzee (Fig. 4c), again, in agreement with tract-tracing data and tractography data in macaques [17,20].

Streamlines from the arcuate in chimpanzees reached inferior frontal gyrus; temporal connectivity was restricted to the posterior components of the STG and STS (Fig. S1a). In contrast, human arcuate streamlines reached all three lateral temporal gyri and extended further anteriorly in the temporal lobe (Fig. S1a). Macaque arcuate streamlines spanned posterior STG to premotor cortex (Fig. S1c), consistent with previous results on arcuate organization in macaques [3,6].

Superior longitudinal fascicle I (SLF I) extended from the superior parietal lobule to the superior frontal gyrus in chimpanzees, humans, and macaques (Fig. 5). Superior longitudinal fascicle III (SLF III) had similar courses for all three species as well: anteriorly reaching inferior frontal gyrus for humans and chimpanzees, the frontal operculum in macaques, and posteriorly, extending to inferior parietal territories (Fig. 5). The second superior longitudinal fascicle (SLF II) ran medially to and in close apposition with SLF III in chimpanzees and macaques (Figs. 5a, 2c), but humans, it was displaced superiorly and is roughly equidistant between SLF I and SLF III (Fig. 5b).

**Figure 5.**
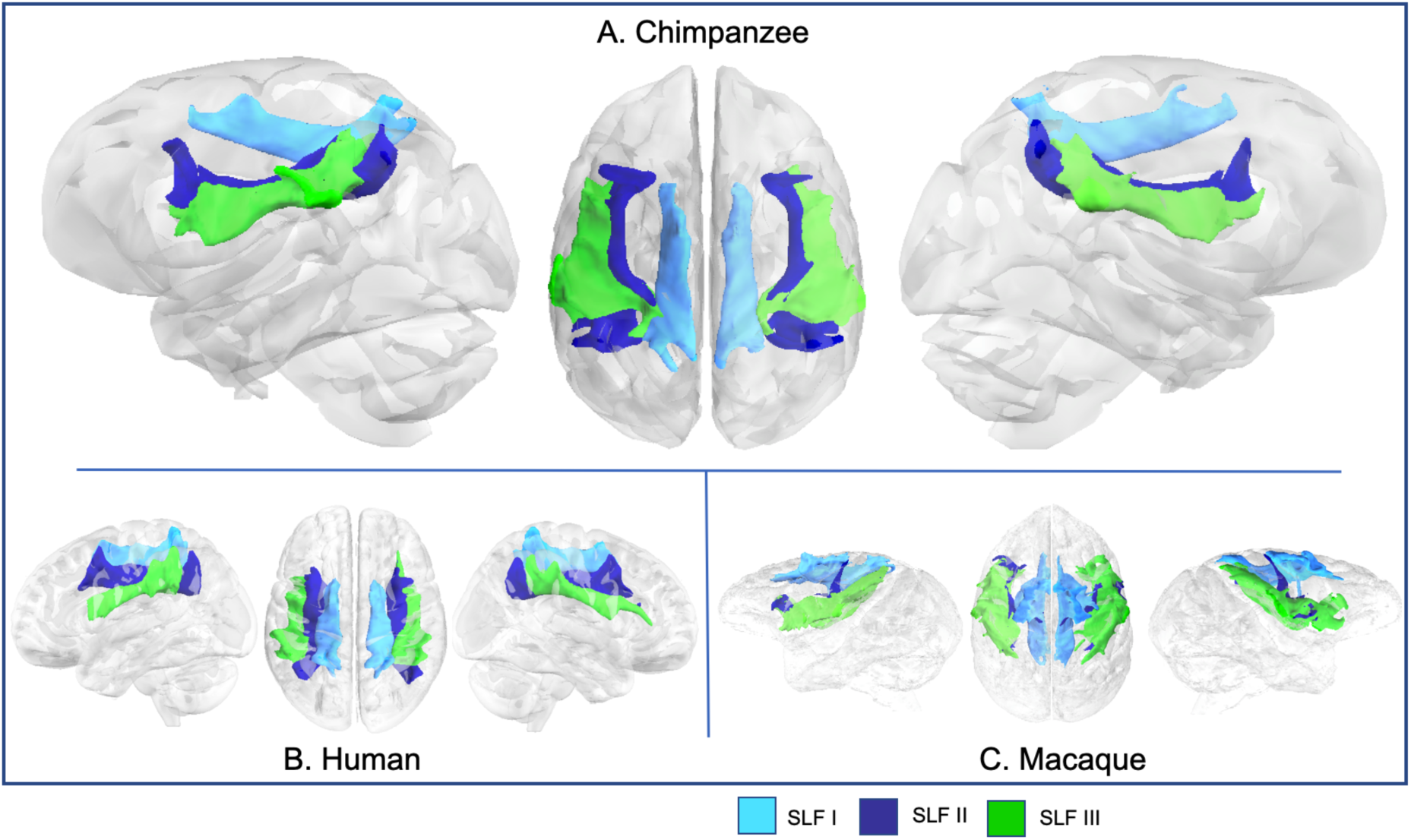
SLFs I, II, and III in chimpanzee (a), human (b), and rhesus macaque (c).

SLF I reached similar areas all three species: superior parietal areas, superior frontal gyrus (SFG), and the superior parietal lobule (Fig. S1b). For SLF II, chimpanzees were similar to humans, with streamlines concentrated in middle frontal gyrus (MFG) and angular gyrus (Fig. S1c), although chimpanzee SLF II also reached the superior parietal lobule. In macaques, SLF II anterior projections extend from the frontal operculum to the upper bank of the arcuate sulcus (Fig. S1c). SLF III projections were also largely consistent across species, with streamlines concentrated in the inferior parietal lobule and inferior frontal gyrus/frontal operculum (Fig. S1d). In chimpanzees, these projections reached the supramarginal gyrus and extended into superior parietal areas. Humans had a similar pattern of posterior projections; anterior projections reached IFG at the level of the pars triangularis (Fig. S1d).

### Temporal association tracts

The middle longitudinal fascicle (MdLF) in chimpanzees spanned the length of STG, reaching from the posterior terminus of the Sylvian fissure to the temporopolar region of anterior STG (Fig 6a). Macaque MdLF also followed this pattern (Fig. 6c), while human MdLF extended further towards the occipital lobe (Fig. 6b). Surface projections show chimpanzee, macaque, and human MdLF connects to the length of STG and most of STS (Fig. S2a). Human MdLF streamlines were more expansive, reaching the superior parietal lobule and the lateral occipital lobe (Fig. S2a).

**Figure 6.**
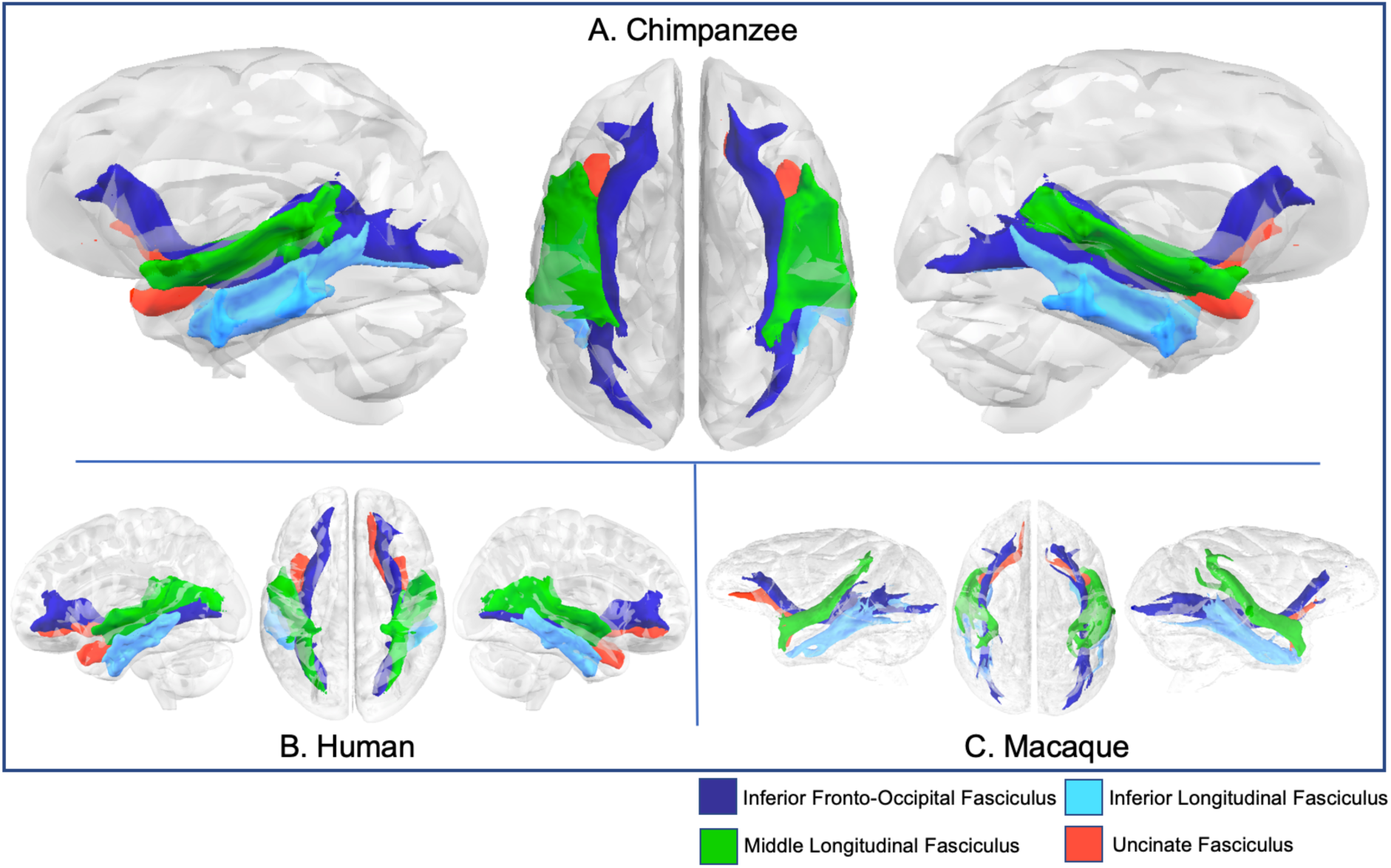
Major fasciculi that course through the temporal lobe (IFOF, ILF, MdLF, and UF) in chimpanzee (a), human (b), and rhesus macaque (c).

In chimpanzees, the inferior longitudinal fasciculus (ILF) coursed through the MTG and ITG, spanning from temporopolar regions to posterior STG and extending toward inferior lateral occipital cortex (Fig 6a), similar as described previously [21]. Human ILFs were similar; macaque ILF coursed through ITG (Figs. 6b, 6c). Streamlines from chimpanzee ILF extended the length of MTG, ITS, and ITG, reaching the angular gyrus posteriorly (Fig. S2b). Human ILF reached the same areas, and macaque ILF reached homologous landmarks - ITG, STS, the upper bank of the STS, and reached posteriorly to the margins of the intraparietal sulcus and the lunate sulcus (Fig. S2b).

The inferior fronto-occipital fasciculus (IFOF) in chimpanzees extended from the occipital lobe, through the temporal lobe, medial to the ILF and MdLF, and into the prefrontal cortex via the extreme/external capsule, coursing superiorly to the uncinate fascicle (Fig. 6a). Humans, compared to chimpanzees, had a similar organization, however the IFOF was slightly inferior to the MdLF, particularly in the posterior temporal lobe (Fig. 6b). Conventionally, IFOF is not a recognized tract in macaques, but recently various authors [22] have proposed that the IFOF is a valid bundle in this species. For the sake of comparison, we reconstructed the macaque tract here using a comparable protocol with the chimpanzee and human, and found a bundle with a similar anatomy coursing medially to MdLF and ILF (Fig. 6c). In all three species the tract extends into prefrontal cortex, where it splits into superior and inferior terminations (Fig. 6).

These patterns are also discernible in surface projections, which show connectivity to both IFG (macaque: frontal operculum) and the frontal pole in all three species (Fig. S2c). Chimpanzee IFOF also projects to angular gyrus and the superior parietal lobule, and occipital cortex (Fig. S2c). Human IFOF streamlines stretch further dorsally in prefrontal cortex and further inferiorly, into fusiform gyrus and ventral occipital cortex. In the macaque, IFOF streamlines are concentrated in STS, STG, in addition to frontopolar and ventral prefrontal areas (Fig. S2c).

In chimpanzees, the uncinate fascicle (UF) extended from the temporopolar region of the STG to inferior frontal cortex, passing through the extreme/external capsule in close apposition and just inferior to the IFOF (Fig. 6a). Human and macaque UF showed comparable anatomy (Fig. 6b & c), but differences were apparent in the surface projections - chimpanzee uncinate reached frontopolar, orbitofrontal, medial prefrontal, and superior temporopolar cortices (Fig. S2d). Human uncinate projections were similar, while macaque uncinate projected to STG, and orbitofrontal cortex (Fig. S2d).

### Limbic tracts

The cingulum bundle was reconstructed by combining the results from three segments - dorsal, prefrontal, and temporal, and had comparable trajectories in all three species - extending from the parahippocampal gyrus, through medial posterior temporal lobe, coursing rostrocaudally superior to the corpus callosum, and terminating in medial prefrontal cortex (Fig. 7). Dorsal prefrontal projections were discernible in all three species, and posterior parietal projections were also visible in humans (Fig. 7b) and macaques (Fig. 7c). Organization of the fornix was also similar in chimpanzees, humans, and macaques - extending from the medial temporal lobe just superior to the cingulum, to the mammillary bodies and hypothalamus (Fig. 7). Surface projection patterns for limbic tracts in chimpanzees were comparable to human and macaque results (Figs. S3a-d).

**Figure 7.**
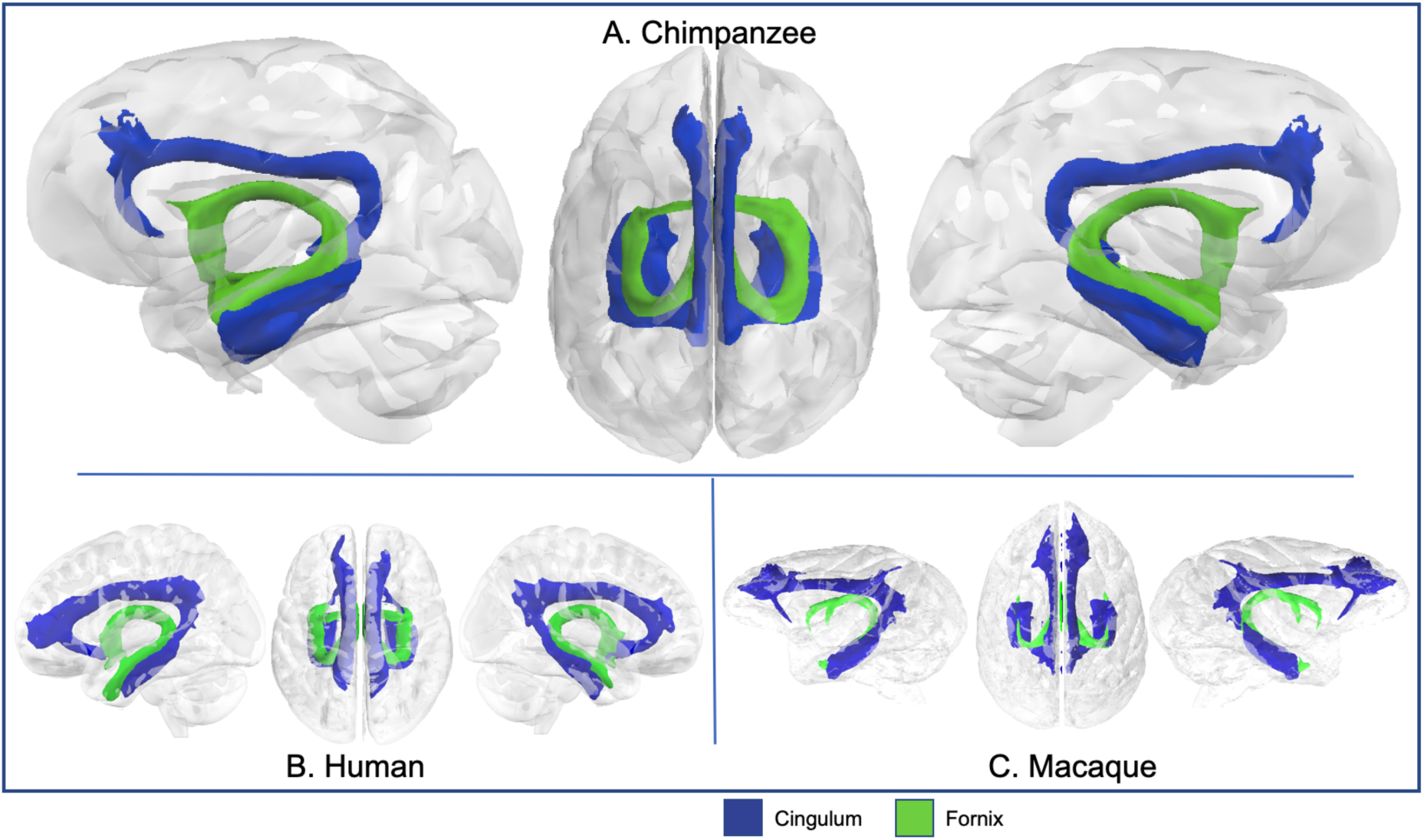
Cingulum and fornix in chimpanzee (a), human (b), and rhesus macaque (c).

### Short tracts

The frontal aslant connects ventrolateral prefrontal cortex with dorsal frontal cortex. In chimpanzees, this tract runs from the inferior frontal gyrus to the superior frontal gyrus, superior to the anterior commissure (Fig. 8a). Human frontal aslant trajectory is similar, but with a straighter fiber bundle owing to the more vertical shape of the human frontal cortex (Fig. 8b). Chimpanzee and human frontal aslant streamlines were concentrated in superior, and inferior frontal gyri; in humans the latter was concentrated in the pars opercularis and triangularis (Fig. S4a). Surface projections in macaque were concentrated in frontal operculum, along both banks of the arcuate sulcus, and dorsal prefrontal cortex (Fig. S4a).

**Figure 8.**
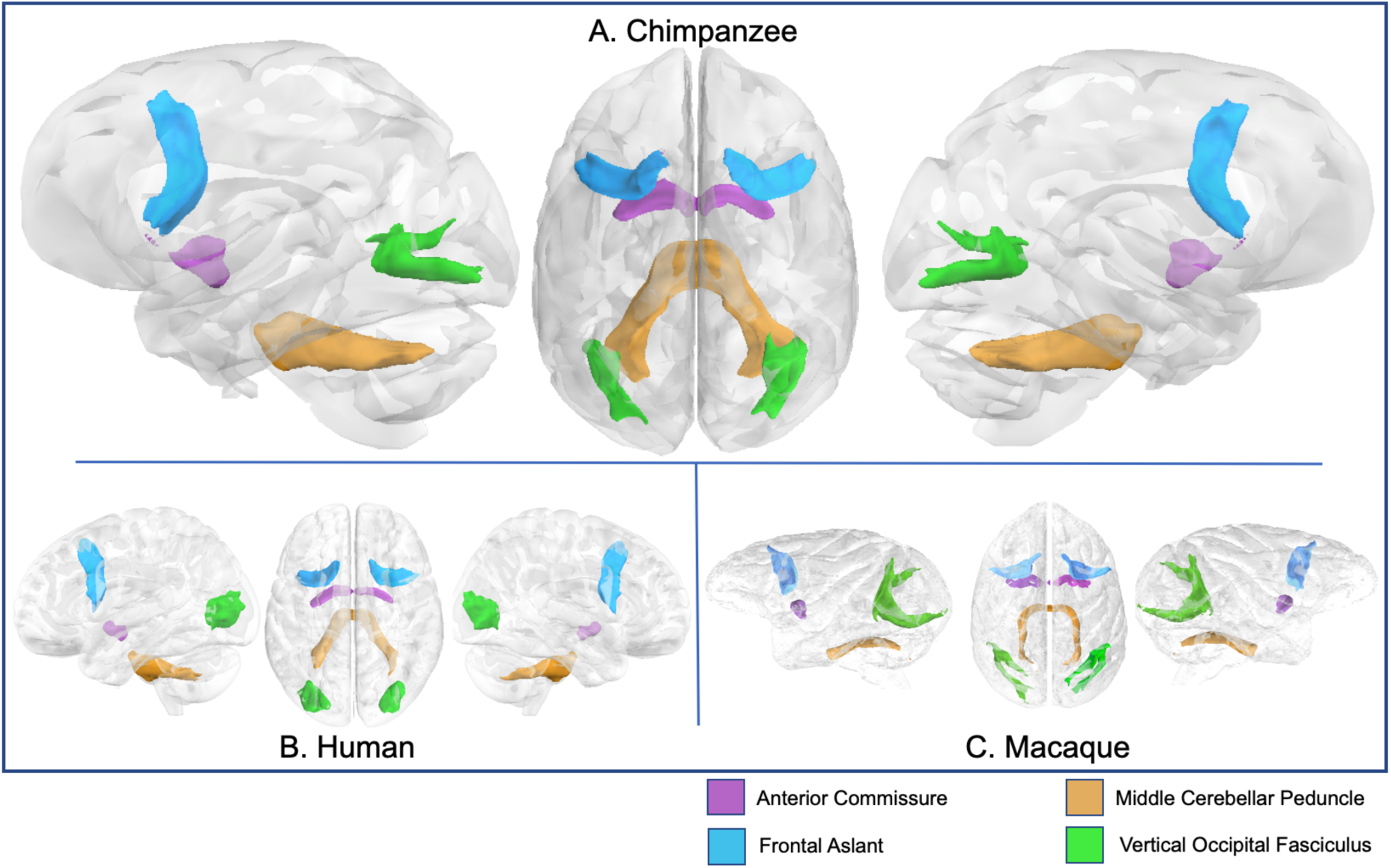
Anterior commissure, frontal aslant, MCP, and VOF in chimpanzee (a), human (b), and rhesus macaque (c).

The vertical occipital fasciculus (VOF) connects dorsal and ventral surfaces of the occipital lobe. In chimpanzees, this bundle arches between these territories, medial to the lunate sulcus (Fig. 8a). In macaques, this bundle is similar in shape but relatively larger, likely due to the larger proportion of space that the occipital lobe takes up in the macaque brain (Fig. 8c). Human VOF was proportionally smaller and much more compact, angled more medially, perhaps due to the rotation of visual cortex to the medial surface of the brain in humans (Fig. 8b). Chimpanzee VOF streamlines projected throughout occipital cortex, to lateral lunate sulcus and inferior to the inferior occipital sulcus (Fig. S4b). Macaque VOF projects along the lunate sulcus, while human VOF streamlines reached most of lateral occipital lobe, including inferior, middle, and superior occipital gyri (Fig. S4b).

### Corticospinal and Somatosensory Pathways

The corticospinal and somatosensory pathways (CSP) send projections from the motor and somatosensory cortices to the spinal cord. The CSP tractogram in chimpanzees appeared similar to humans and macaques (Fig. 9), and all three species showed tracts reaching precentral and postcentral gyri (Fig. S3e).

**Figure 9.**
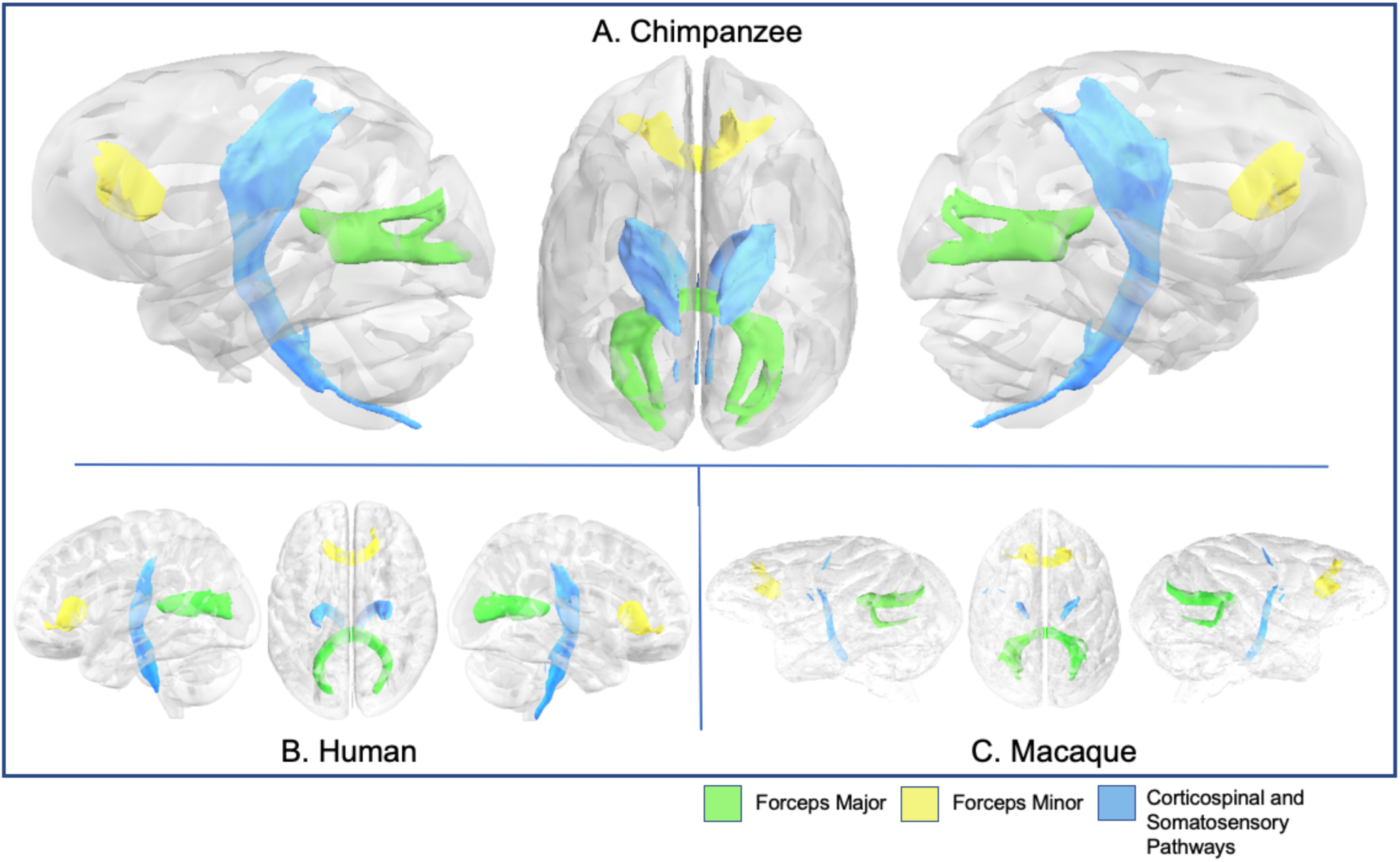
CSP, forceps major and minor in chimpanzee (a), human (b), and rhesus macaque (c).

### Interhemispheric tracts

The anterior commissure connects ventral and anterior temporal cortices of both hemispheres, including the amygdalae. The organization of the anterior commissure was comparable in all three species (Fig. 8, S4c). The middle cerebellar peduncles are a collection of fiber tracts which arise in the pontine nuclei and project to the opposite cerebellar hemisphere; these appeared similar in chimpanzees, humans, and macaques (Fig. 8).

The forceps major and minor are components of the corpus callosum (passing through the splenium and the genu, respectively). Forceps major tractograms in chimpanzees appeared to have a superior and inferior branching as they reached the occipital lobe in chimpanzees and macaques (Figs. 9a, 9c), unlike humans, which either had one branch, or the branches are compressed together due to the different morphology of the occipital lobe in humans (Fig. 9b). Forceps major streamlines reached the medial occipital lobes in all three species, particularly concentrated in the calcarine and parieto-occipital sulci (Fig. S4d). Forceps minor bundles appeared similar in all three species (Fig. 9); their streamlines reached mediodorsal prefrontal cortex, SFG, MFG, and IFG in chimpanzees and humans, and comparable territories (frontal operculum, frontal pole, and medial prefrontal cortex) in macaques (Fig. S4e).

### Thalamic projections

Anterior, superior, and posterior thalamic radiations project from the thalamus to prefrontal, frontal, and occipital cortices, respectively. Compared with macaques, chimpanzees and humans had more robust, vertically expanded tracts (Fig. 10). Perhaps due to the pattern of prefrontal expansion in human cortex, chimpanzee anterior thalamic radiations appeared to be rotated slightly dorsally compared with humans. However, surface projection results demonstrated that this radiation reached SFG, MFG, and IFG in both species (Fig. S5a). In macaques, these streamlines spanned frontal operculum to principal and arcuate sulci (Fig. S5a). The superior thalamic radiations of all species reached precentral gyri and, in humans and chimpanzees, the posterior portion of the superior frontal gyrus (Fig. S5b). Posterior thalamic radiations in chimpanzees and macaques were concentrated in the occipital lobe, from the posterior end of the calcarine fissure to the lunate sulcus, while in humans it reached to all major occipital gyri (Fig. S5c).

**Figure 10.**
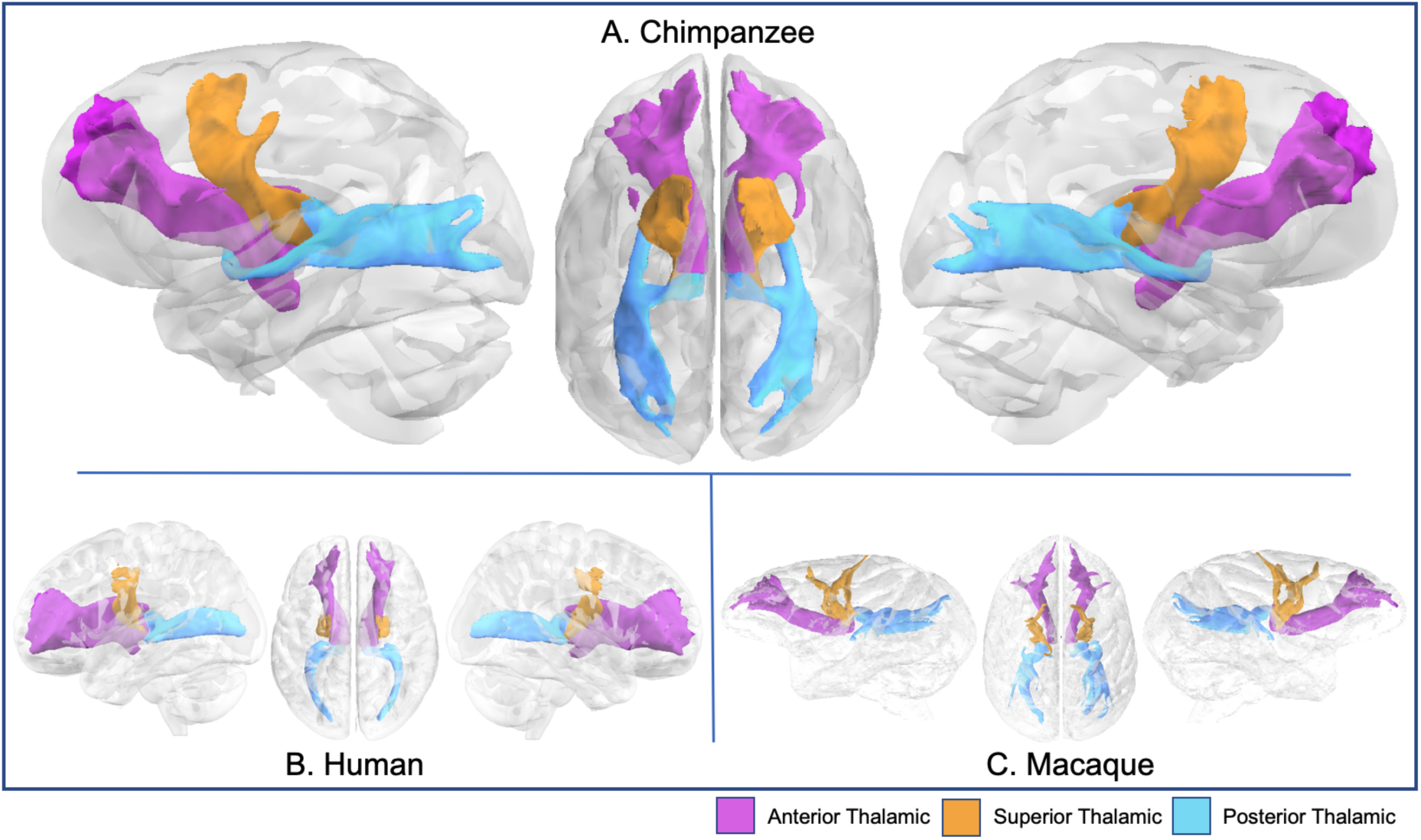
Anterior, superior, and posterior thalamic radiations in chimpanzee (a), human (b), and rhesus macaque (c).

Optic and acoustic radiations connect the small geniculate nuclei (middle geniculate nuclei for acoustic; lateral for optic) to the primary sensory cortices. Chimpanzee optic and acoustic radiation tractograms were comparable to humans and macaques (Fig. 11). As expected, optic radiation streamlines were mainly restricted to the calcarine fissure in chimpanzees and humans, and the occipital pole in macaques (Fig. S5d) while acoustic radiation stnesreamli were concentrated in the planum temporale and posterior STG in chimpanzees, humans, and macaques (Fig. S5e).

**Figure 11.**
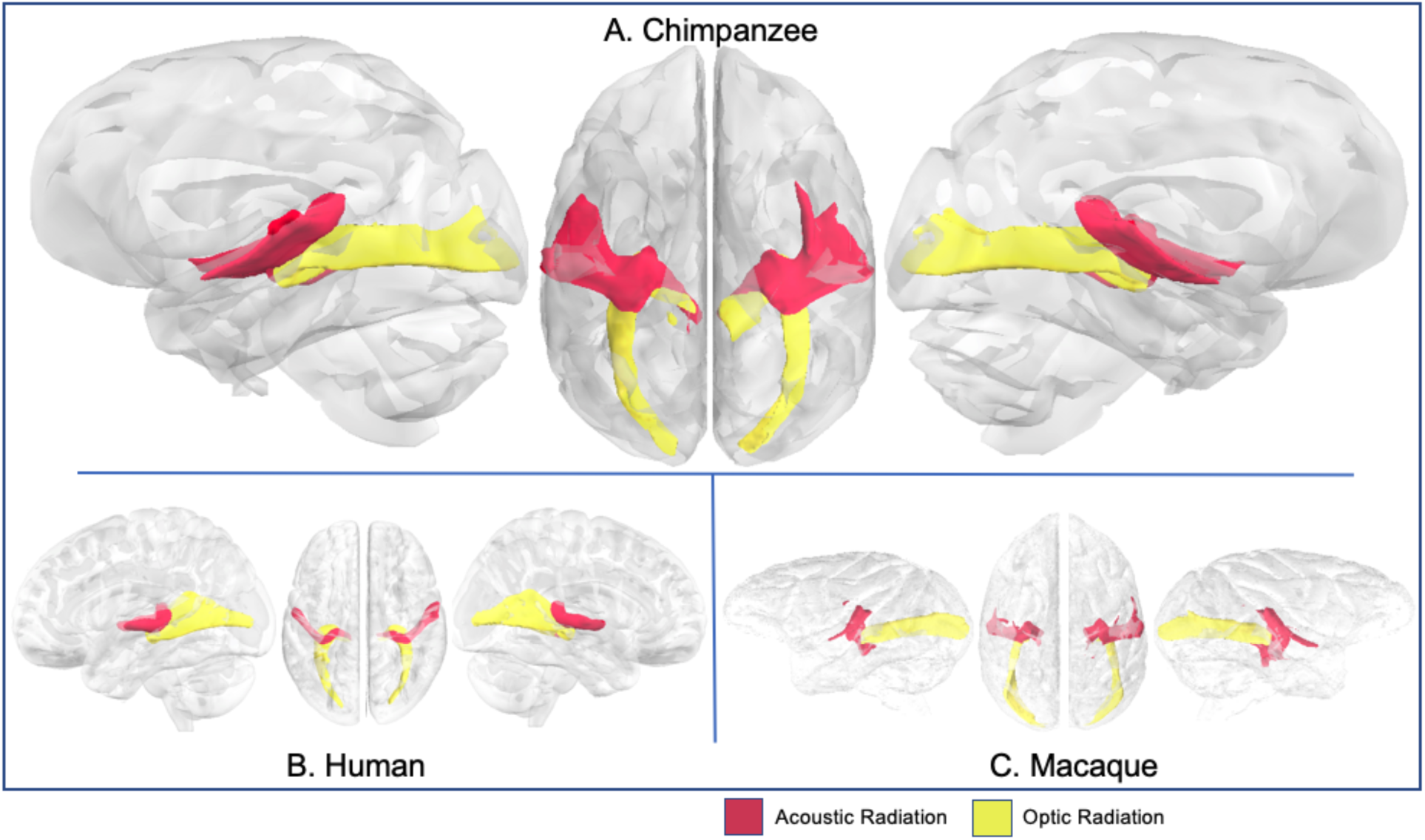
Optic and acoustic radiations in chimpanzee (a), human (b), and rhesus macaque (c).

### Deterministic tractography

To validate our results with an alternative tractography algorithm, we performed deterministic tractography in 5 individual chimpanzees in a subset of tracts (Fig. S6). The course of the tracts is highly similar to those described above and robust across individuals.

## Discussion

We have presented a library of recipes to identify the major white matter tracts in the chimpanzee brain. Similar libraries, based on a similar set of recipes, are available in the human and macaque brain [8,11] and we have presented the results of all three species here for comparison. Creating such a library for the chimpanzee, one of our closest animal relatives, allows direct quantitative investigations on general principles of primate brain organization as well as on specializations of great apes.

By using diffusion MRI tractography to identify the white matter tracts we capitalize on two major advantages of the technique. First, the non-destructive and non-invasive nature of MRI means that it can be used across a large range of species [23]. Traditional techniques such as tract tracing are not feasible in great apes for both practical and ethical reasons. Although diffusion MRI tractography is not without its drawbacks (see below), the ability to use the same technique in all species provides a unique opportunity for comparative research. Additionally, the digital nature of the data means that they are easily shared -- different labs can exchange their protocols easily and use them for their tractography algorithm of choice, including deterministic tractography (fig. S6). This facilitates the use of common terminology, offers the chance for others to build upon and improve our recipes, and will aid in solving disputes of tract identifications. To this end, the current library is compatible with the XTRACT tool of the FSL software [11], facilitating easy standardization and exchange.

In the next sections, we will discuss how the tracts identified by our recipes can enrich our understanding of white matter evolution in humans and chimpanzees, and explore some possible functional implications for these similarities and differences. However, in-depth quantitative investigations comparing white matter architecture of the chimpanzee with that of other species, focusing for instance on the evolution of hemispheric lateralization [24] and identification of grey matter homologies, cf. [25], will be reserved for future communications.

### Implications for Language Evolution

The arcuate fasciculus, IFOF, MdLF, and uncinate have all been implicated in language abilities in humans, and are therefore of interest to comparative neuroanatomists studying the evolution of language and language-like abilities of primates [26]. The arcuate, in particular, is one of the most rigorously examined tracts, and is one of the few fasciculi previously examined in chimpanzees [6]. Human arcuate has extensive connections linking STG, MTG, and ITG in the temporal lobe to Broca’s area (BA 45 and BA 44), ventral premotor cortex, and middle frontal gyrus in prefrontal cortex [27,28]. Some workers recognize an additional component, the posterior segment, connecting temporal cortex to the inferior parietal lobule [29]. By integrating language comprehension areas with speech production areas, the left arcuate, also known as the phonological pathway [30], facilitates language, particularly speech production, in humans [31,32]; [33,34] but see [35].The right arcuate is implicated in processing music [36] and pitch [37].

We find chimpanzee arcuate to have a similar organization to humans in the dorsal component, but with a less extensive temporal projection, restricted to STG, which is consistent with previous studies [6,38]. Compared with humans, the temporal component of the arcuate in chimpanzees is much smaller and does not project beyond the posterior and superior portions of the temporal lobe, similar to what we observed in our macaque dataset (Figs. 1, 9a). By contrast, prefrontal connectivity in chimpanzees was similar to humans, with robust tracts reaching further anteriorly than macaques, into the pars triangularis of the IFG. Human facility with language, including lexical retrieval, semantic richness, complex syntax, and precise articulation, is likely related to the expansion of this tract into the middle and inferior portions of the temporal lobe and the inferior frontal gyrus in prefrontal cortex. The ability of chimpanzees and bonobos to acquire some semantic comprehension and manual language production, e.g. [39–41], suggests that the connections between temporal and prefrontal areas that we see in the chimpanzee but not the macaque - specifically, those in prefrontal areas - enable these cognitive abilities.

The “dual streams” model of human language disambiguates the arcuate “dorsal” pathway from the IFOF “ventral” pathway [42,43]. In macaques, the extreme and external capsule fiber bundles have been described [17], but many workers argue that this tract does not reach the occipital cortex to form a true IFOF in this species, instead terminating in the STS [3,13,44]. However, *ex vivo* diffusion tractography [22,45] and fiber dissection [46] studies have identified occipital projections from the extreme/external capsule in macaques consistent with IFOF, and similar results have been shown for the marmoset monkey [47] and vervet monkey [48]. These results can be reconciled with the tracer data if one assumes that the tractography and blunt dissection results both pick up multi-synaptic pathways.

In our dataset, chimpanzee IFOF reached areas 45 but not 44, based on the anatomical locations of these homologues as previously described [49,50]. Human and macaque IFOF also reached area 45 (pars triangularis in humans) but not 44 (pars orbitalis in humans), consistent with previous tractography studies in these species [51], suggesting that the organization of anterior IFOF may have evolved in the ancestor of Old World anthropoids. Posterior connectivity projections had shared features between humans and chimpanzees, reaching inferior parietal and lateral and inferior occipital areas. In macaques, IFOF reached the upper bank of the STS, but did not extend into inferior parietal or occipital areas beyond the lunate sulcus. Together, our data suggest that IFOF connectivity may be a necessary but not sufficient condition for human-like language abilities.

### Differences Due to Morphometric Changes

Of the tracts we examined, several appeared to have important differences between species on visual inspection; however closer examination demonstrates that changes in the morphology of the tract in question were due to changes in the overall size and shape of the brain and not changes in connectivity. For example, the vertical occipital fasciculus appears quite divergent between species (Fig. 5). This is a consequence of the cortical expansion that has occurred in the hominin lineage; posterior and inward displacement of primary visual cortex in chimpanzees and to an even greater extent in humans [52]. Chimpanzees, like macaques, have primary visual cortex located on the lateral surface of the occipital lobe, delineated by the lunate sulcus, but in chimpanzees, the area bounded by the lunate takes up a smaller proportion of the cortex when viewed from the side, as a result of parietal and temporal association cortex expansion. Human V1 is fully displaced medially and inferiorly, and is not bounded by the lunate. The endpoints of the VOF are therefore compressed vertically relative to the rest of the brain in chimpanzees and humans.

The uncinate fascicle in chimpanzees and humans showed a bend in its trajectory of approximately 45 degrees as it moves between the temporal and prefrontal cortices; but in macaques we see a gentler curve of about 90 degrees (Fig. 3). Again, the course of this tract was similar in all three species; connecting the superior anterior temporal lobe to ventral prefrontal areas. Here, longitudinal expansions to the temporal lobe of chimpanzees and humans [26] are responsible for the change in angle to this tract.

An examination of the thalamic radiations across species (Fig. 7) offers a viewpoint on the overall expansion trends of chimpanzee and human cortex. Compared with macaques, the orientation of the STR and PTR are at a greater angle in the chimpanzee. When you compare the human to the chimpanzees, we see that there is a greater angle between the ATR and STR in humans. This may reflect patterns of expansion in parietal and frontal association territories - particularly, the overall increase in globularity in the human brain resulting from the expansion of these areas in the hominin lineage [53,54].

### Considerations and Limitations

Our chimpanzee dataset was originally collected as part of a study on female primate aging prior to 2012, and for that reason, consisted of female individuals only. Thus, our chimpanzee and human datasets are reasonably age-matched, but not sex-matched. However, reported sex differences in fasciculi between adult male and female humans are small and limited to the left SLF [55]. Group sex differences found in human brain morphometry have reproducibility problems [56–58], and to the degree to which they are reproducible, are not categorical differences but rather differences of mean values, with high degrees of population overlap (e.g., [59]; for reviews, see [60,61]). Crucially, sex differences are often a matter of total volume and not anatomical organization of fascicular projections, and therefore we would not expect our anatomical descriptions to change if male chimpanzee scans were added to the dataset. Despite this, it would be preferable to include male chimpanzees in our analysis. It is possible there are greater sex differences in chimpanzees than humans, in which case it would be important to capture male neuroanatomy in our characterization of “the chimpanzee”. Future studies will expand the dataset to include male chimpanzees that were scanned for a different study, and our publicly available dataset will be updated as improvements to the dataset are implemented.

Diffusion MRI tractography is a relatively new tool for comparative neuroscience. Although it has proven to be replicable and has obvious advantages of its wide applicability, it has not been without its criticisms when directly compared to more traditional neuroscientific methods. Promisingly, studies comparing tract-tracing with diffusion tractography in *ex vivo* macaques have found comparable results[15,62–64]. Moreover, high angular resolution data such as used in the present investigation has been shown to perform well on difficult-to-reconstruct tracts like the acoustic radiation[65], and multifiber algorithms increase sensitivity[10]. However, a number of issues remain, which we will discuss in detail below.

Tractography can produce false positives, and the best way to mitigate this is the use of strong anatomical priors [66]. Here, we developed standardized protocols based on strong anatomical knowledge from other species, including the macaque in which a wealth of tracer data is available. Additionally, alternative definitions of tracts can be added to the recipe library to arbitrate putative false positives. Correcting for false positives can open up the opposite problem of false negatives; to guard against this, these data can be compared with results from data-driven methods that identify tracts without using anatomical priors. Our recent study using this approach [67] found results broadly consistent with those obtained here. The largest difference we observe is that SLFs are difficult to reconstruct in the chimpanzee without strong anatomical priors.

Another issue for tractography is in reaching the grey matter. Superficial white matter and a bias in tracking towards gyral crowns can impair accurate identifications of connections leaving the white matter and entering grey matter [14,68]. These problems do not impact our main results, where we solely identify the course of major fibers through white matter. To mitigate gyral bias in our surface projections, we track from the gray matter towards the whole-brain white matter instead of vice versa, avoiding the problem of passing through a gyral bottleneck and having to reconstruct fanning fibers within the gyrus, and then multiply the resulting matrix with the tractogram of each tract, cf. [8].

DTI tractography in comparative anatomical studies must also reckon with size differences and scan resolution differences, which can impact results. Chimpanzee and human diffusion scans had 1.5 mm isotropic resolution, however, chimpanzee brains are approximately one third the size of human brains. Higher resolutions are achievable with post mortem tissue, and our *ex vivo* macaque scans had very good resolution (0.6 mm^3^). Going forward, the development of larger *ex vivo* datasets for all species will permit even more precise comparisons between species.

Despite these limitations, diffusion tractography offers several indispensable benefits for comparative neuroanatomical research. Very few methodologies exist that can be used in humans, chimpanzees, and macaques. dMRI is noninvasive and can be used on *ex vivo* tissue for long time periods to produce high quality scans, especially valuable for researchers who are interested in smaller-brained species. Although the quality of *ex vivo* scans is highly dependent on the time interval between death and fixation and in general suffer from a lower SNR [69], the availability of dedicated sequences coupled will well-established protocols for sample preparation

[70] now make this approach feasible (e.g., [21,22,71]). It should be stressed that diffusion tractography should not be understood as precisely the same as tract-tracing data, which reveal specific anatomical connections, but rather, as a method for characterizing white matter bundles on a larger scale, their spatial relationship with one another, and with cortex.

### Conclusions

In conclusion, these data indicate that 1) prefrontal projections of the chimpanzee arcuate are similar to humans and different from macaques; 2) temporal projections of the chimpanzee arcuate are similar to macaques and different from humans; and 3) posterior connections of the IFOF have been modified in both human and chimpanzee lineages.

We have come a long way since the first comparative anatomical studies of chimpanzee brains identified the presence of the “hippocampus minor” as the distinguishing feature of humans [72]. In this new era, more complex and detailed maps at multiple levels are illuminating the similarities and differences between humans and chimpanzees, our closest relatives, and macaques, commonly used as animal models for human diseases and therefore important to human health. Comparative neuroimaging facilitates phylogenetic analysis which can pinpoint the areas of human brains that are critical for human cognitive abilities like language, conceptual processing, tool use, and sophisticated social cognition [23,73].

This chimpanzee white matter atlas is presented as a resource paper and a starting point for future quantitative and comparative analysis. Future work will take advantage of this atlas and accompanying tract recipes to reconstruct tracts in different chimpanzee populations, to use as a reference to create tractography recipes for other primate species, and to more finely probe the neuroanatomical adaptations unique to humans.

## Materials and Methods

### Chimpanzee data

Chimpanzee (Pan troglodytes; n=29, 28 ± 17 yrs, all female) MRI scans were obtained from a data archive of scans acquired prior to the 2015 implementation of U.S. Fish and Wildlife Service and National Institutes of Health regulations governing research with chimpanzees. All the scans reported in this publication were collected as part of a grant to study aging in female primates; all scans were completed by the end of 2012 and have been used in previous studies, e.g. [74,75]. All chimpanzees were housed at the Yerkes National Primate Research Center (YNPRC) in Atlanta, Georgia, USA. All procedures were carried out in accordance with protocols approved by the YNPRC and the Emory University Institutional Animal Care and Use Committee (IACUC approval #YER-2001206).

Chimpanzee subjects were immobilized with ketamine injections (2–6 mg/kg, i.m.), and then anesthetized with an intravenous propofol drip (10 mg/kg/h) prior to scanning, following standard YNPRC veterinary procedures. Subjects remained sedated for both the duration of the scans and the time necessary for transport between their home cage and the scanner location. After scanning, primates were housed in a single cage for 6–12 h to recover from the effects of anesthesia before being returned to their home cage and cage mates. Veterinary and research staff evaluated the well-being (activity and food intake) of chimpanzees twice daily after the scan for possible post-anesthesia distress.

Anatomical MRI and diffusion MRI scans were acquired in a Siemens 3T Trio scanner (Siemens Medical System, Malvern, PA, USA). A standard circularly polarized birdcage coil was to accommodate the large chimpanzee jaw, which does not fit in the standard phase-array coil designed for human subjects. Diffusion-weighted MRI data were collected with a single-shot, spin-echo echo-planar imaging (EPI) sequence. A dual spin-echo technique combined with bipolar gradients was used to minimize eddy-current effects. Parameters were as follows: 41 slices were scanned at a voxel size of 1.8 x 1.8 x 1.8 mm, TR/TE: 5900 ms/86 ms, matrix size: 72×128. Two diffusion-weighted images were acquired for each of 60 diffusion directions, each with one of the possible left–right phase-encoding directions and eight averages, allowing for correction of susceptibility-related distortion [76]. For each average of diffusion-weighted images, six images without diffusion weighting (b = 0 s/mm^2^) were also acquired with matching imaging parameters. High-resolution T1-weighted MRI images were acquired with a 3D magnetization-prepared rapid gradient-echo (MP-RAGE) sequence for all subjects. T2 images were previously acquired [77] using parameters similar to a contemporaneous study on humans [78].

Data were analyzed using tools from the FSL software library of the Oxford Center for Functional Magnetic Resonance Imaging of the Brain (FMRIB; www.fmrib.ox.ac.uk/fsl/;[79]. T1-weighted images were skull-stripped using BET, with some manual correction [80] in the posterior occipital lobe. FSL’s *eddy_correct [81]* and *topup* [82] implemented in Matlab (Matlab7, Mathworks, Needham, MA) were used to correct eddy current and susceptibility distortion in the diffusion data. Diffusion-weighted images were processed using FMRIB’s Diffusion Toolbox, to fit diffusion tensors, estimate the mean diffusivity and fractional anisotropy, and fit with a voxel-wise model of diffusion orientations using bedpostX, using a crossing fiber model with three fiber directions [10]. In addition to standard T1w/T2w and diffusion data preprocessing, a modified version of the Human Connectome Project (HCP) minimal preprocessing pipeline [83] was used to create cortical surfaces and registrations to a population-specific chimpanzee template.

Template generation for chimpanzees has been previously described [84,85]; briefly: the PreFreeSurfer pipeline was used to align the T1w and T2w volumes of 29 individual chimpanzees to native anterior commissure-posterior commissure space. Brain extraction, cross-modal registration, bias field correction, and nonlinear volume registration to atlas space were performed using FSL (FMRIB Software Library, University of Oxford [79]), and cortical thickness was computed using the FreeSurfer mris_make_surfaces command. The PostFreeSurfer pipeline was used to produce a high-resolution 164k surface mesh (∽164,000 vertices per hemisphere), as well as a lower-resolution mesh (32k for chimpanzee).

### Human and macaque data

Three post mortem macaque brain scans (*Macaca mulatta*, n=3; two male, age range 4-14 years) were acquired using a 7T magnet with an Agilent DirectDrive console (Agilent Technologies, Santa Clara, CA, USA). A 2D diffusion-weighted spin-echo protocol was implemented with a single line readout (DW-SEMS, TE/TR: 25 ms/10 s; matrix size: 128 x 128; resolution: 0.6 x 0.6 mm; number of slices: 128; slice thickness: 0.6 mm). Nine non-diffusion-weighted (b=0 s/mm^2^) and 131 diffusion-weighted (b=4000 s/mm^2^) volumes were acquired with diffusion directions distributed over the whole sphere. Before scanning, brains were soaked in PBS; they were placed in fomblin for the duration of the scan. The b=0 images were averaged and spatial signal inhomogeneities were restored. Although diffusion anisotropy is largely preserved, *ex vivo* tissue usually has reduced diffusivity, partially due to fixation [69] and lower temperature (room temperature instead of body temperature), necessitating the use of larger b-values to achieve equivalent diffusion contrast to *in vivo* data; this was achieved here by increasing the diffusion coefficient from b = 1000 to 4000 s/mm^2^ [86,87]. Diffusion-weighted images were processed using the same method as chimpanzees, described above. The cortical surface of one macaque with high quality structural MRI was reconstructed using a modified version of the HCP pipeline. A nonlinear registration between this macaque and the other macaques was obtained using FLS’s nonlinear registration tool FNIRT [88] based on the T1w scans. The surface was then warped to the other macaque brains to enable using the same surface model in all three macaques and then transformed to F99 standard space [89].

Nineteen human subjects (10 female, 2 ages 22-25; 8 ages 26-30; 9 ages 31-35) were selected from the *in vivo* diffusion MRI data provided by the Human Connectome Project (HCP), WU-Minn Consortium (Principal Investigators: David Van Essen and Kamil Ugurbil; 1U54MH091657) funded by the 16 NIH Institutes and Centers that support the NIH Blueprint for Neuroscience Research; and by the McDonnell Center for Systems Neuroscience at Washington University [90]. Minimally preprocessed datasets from the Q2 public data release were used. Data acquisition and preprocessing methods have been previously described [83,90,91]. In brief, diffusion data were processed in an analogous manner to the chimpanzee and macaque data, resulting in a voxel-wise model of diffusion orientations yielded by FSL’s bedpost [92] while T1 and T2-weighted data were used for surface reconstruction similar to that described above for the chimpanzee.

### Tractography

Probabilistic diffusion tractography was performed using FSL’s probtrackx2 [10]. Region-of-interest (ROI) masks were manually drawn in template volume space in all three species. Recipes consisting of seed, waypoint (target), exclusion, and/or stop masks for each white matter tract of interest were developed in chimpanzees following the approach previously established for the human and macaque [8,11]. Details of these protocols follow in the next section. Masks for tractography were warped to subject space for tractography, which was run in two directions (seed to target and target to seed). Tractography parameters were set as follows: Each seed voxel was sampled 10,000 times, with a curvature threshold of 0.2, step length of 0.5 mm, and a number of steps of 3200. The resulting tractograms were normalized by dividing by the waypoint number and warped back to template space. The normalized tractograms were then averaged and thresholded at 0.0005 across all individuals to create a single group tractogram for each tract. For comparison, all tracts were also reconstructed in the human MNI152 standard space and in macaque F99 [93]. Tractography was performed for both species analogously to the chimpanzee with the same parameters, with the exception of step length which was changed to 0.25 for macaques. Results were visualized with Matlab (MATLAB and Statistics Toolbox Release 2012b, The MathWorks, Inc., Natick, Massachusetts, United States). Deterministic tractography was run on individual chimpanzees using DSI Studio March 10, 2020 build (http://dsi-studio.labsolver.org; [94]).

### Tractography recipes

Descriptions of chimpanzee protocols are described below. Unless noted otherwise, human and macaque protocols were comparable, and all protocols included a bilateral exclusion mask consisting of a sagittal section isolating tracts within the two hemispheres with the exception of the middle cerebellar peduncle and anterior commissure.

### Dorsal tracts

#### Superior Longitudinal Fasciculi and Arcuate Fasciculus

Superior longitudinal fasciculi (SLFs) and arcuate fasciculus (AF) were reconstructed using a one seed, two target approach, in which the seed ROI was drawn in the center of the tract and a target ROI was drawn at the anterior and posterior ends of the tract (Figs. 1a-c). The chimpanzee arcuate fasciculus was reconstructed with a seed drawn in the white matter medial to the supramarginal gyrus (SMG), around the end of the Sylvian fissure. A temporal target mask was drawn in the white matter encompassing the STG and MTG, and a second target was drawn at the level of the ventral premotor cortex, posterior to the inferior frontal gyrus (IFG) and anterior to the precentral sulcus. A stop mask was drawn in the area encompassing the auditory core to avoid contamination from the acoustic radiation (Fig. 1a).

For SLF I, the seed was drawn slightly anterior to the central sulcus, in the white matter at the base of the superior frontal gyrus (SFG), with one target in the SFG at the level of the precentral gyrus and a posterior target in the superior parietal lobule (Fig. 1b). SLF II involved a seed mask deep to the postcentral sulcus, with target masks in the middle frontal gyrus at the level of the PCG and in the inferior parietal lobule (Fig. 1c). For SLF III, the seed was placed in the white matter of the SMG, with a target in the inferior frontal gyrus and in the posterior SMG, at the posterior terminus of the Sylvian fissure (Fig. 1d). Dorsal tract protocols employed exclusion masks that excluded the basal ganglia, and in the case of SLF I, a portion of the cingulate gyrus, to prevent contamination from the cingulum (Fig. 1b). The placement of posterior target masks was informed by tracking data from humans from Kamali and co-workers [95]. Seed and anterior target placement was guided by a previous DTI study on chimpanzee SLF anatomy [96].

Protocols in humans and macaques were similar with four exceptions: In macaques, the posterior target for the arcuate protocol was located in the axial plane of the STG posterior to the terminus of the Sylvian fissure (Fig. 3a), and additional exclusions of the superior parietal lobule were added to avoid contamination with SLF I (Figs. 3c, d). Finally, arcuate protocols did not require auditory core stop masks, and SLF I protocols did not require cingulate gyrus exclusions in either humans (Figs. 3a, b) or macaques (Figs. 3a, b).

### Temporal tracts

#### Middle Longitudinal Fasciculus

MdLF was reconstructed using a seed in the white matter of the STG, slightly anterior to the central sulcus, and a target also in the STG, just anterior to the posterior terminus of the Sylvian fissure (Fig. 1e). Exclusion masks were placed in the MTG, ITG, and the prefrontal cortex (Fig. 1e), to prevent contamination with IFOF and ILF. For the ILF, seed and target masks were placed in the white matter within the MTG and ITG, at approximately the same coronal section as the MdLF seed and target masks (Fig. 1f). For humans, the target mask was moved posteriorly, extended inferiorly into the occipital lobe and superiorly to reach the level of the angular gyrus. In addition, an axial slice was extended from the superior limit of the mask traversing the inferior parietal lobule (Fig. 2e). These modifications were based on an anatomical study of MdLF which found projections to occipital cortex and the superior parietal lobule in humans [97].

#### Inferior Longitudinal Fasciculus

Seed/target masks and exclusion masks were inverted from the MdLF protocol. The seed mask was located in a coronal section of the MTG/ITG just posterior to the temporal pole, and the target mask was located in an MTG/ITG coronal section just anterior to the level of the terminus of the Sylvian fissure (Fig. 1f). Exclusion masks were placed in coronal sections of the STG, as well as axially and coronally around the hippocampal formation and amygdala, as well as at the cerebellar peduncle (Fig. 1f).

#### Inferior Fronto-Occipital Fasciculus

IFOF recipes involved a large coronal slice in the occipital lobe for the seed and a coronal slice in the prefrontal cortex, just anterior to the genu of the corpus callosum (Fig. 1g). In order to restrict streamlines to those that pass through the extreme/external capsule, a large exclusion mask was drawn as a coronal slice with two lacunae at the extreme/external capsule (Fig. 1g). In macaques, the exclusion mask also included a narrow axial slice at the level of the cingulate in order to eliminate contamination from cingulum fibers (Fig. 3g).

#### Uncinate Fasciculus

Like the IFOF protocol, the uncinate protocol also used a large coronal exclusion mask with lacunae at extreme/external capsule. The seed was placed in the white matter of the STG close to the temporal pole and the target was drawn in the extreme/external capsule (Fig. 1h). A second exclusion mask was placed posterior to the basal ganglia to prevent contamination from longitudinal fibers from the IFOF.

### Limbic tracts

#### Cingulum Bundle

This tract was divided into three separate recipes - cingulum dorsal (CBD), cingulum prefrontal (CBP) and cingulum temporal (CBT). The seed and target for CBD were placed in the cingulate gyrus, at the level of the precuneus and the dorsal part of the genu, respectively (Fig. 1i). The CBP seed mask was placed in the dorsal genu and the target at the ventral terminal point of the genu (Fig. 1j). The CBT recipe involved a seed in the posterior parahippocampal gyrus and one in the anterior parahippocampal gyrus, and two stop masks to avoid picking up occipitotemporal and extreme/external capsule fibers (Fig. 1k).

#### Fornix

The fornix protocol included a seed in the mammillary bodies and a target in the hippocampal formation (Fig. 1l). In addition to the bilateral exclusion mask, a coronal section of occipital lobe was excluded to avoid contamination from posterior fiber bundles (Fig. 1l).

### Short tracts

#### Frontal Aslant

A seed was placed in a parasagittal section of the IFG, at the white matter stem, and the seed in an axial section of the superior frontal gyrus (Fig. 1m). The exclusion mask included a coronal slice just posterior to the seed and target masks, to avoid including streamlines from dorsal fasciculi (Fig. 1m).

#### Vertical Occipital Fasciculus

The vertical occipital fasciculus (VOF) recipe contains a seed in the white matter of the occipital lobe, inferior to the calcarine fissure, and a target in the occipital white matter lateral to the fundus of the calcarine and medial to the fundus of the lunate sulcus (Fig. 1n); because the lunate sulcus in humans is quite small, the target was medial to the lateral sulcus of the occipital lobe in humans (Fig. 2n). An exclusion mask consisting of a coronal section at the level of the posterior temporal lobe was used to avoid the inclusion of longitudinal tracts (Fig. 1n).

### Corticospinal and Somatosensory Pathways (CSP)

A seed was placed in the central medial portion of the pons, with a target encompassing the superior parietal lobule, inferior parietal lobule, and postcentral gyrus (Fig. 1o). Exclusion masks were placed in an axial section, at the level of the midbrain, which excluded streamlines outside of the midbrain/brainstem (Fig. 1o). In the macaque, the target mask covered an area between the central sulcus and the intraparietal sulcus (Fig. 3o).

### Interhemispheric tracts

#### Forceps Major and Minor

A seed mask consisting of a coronal slice of one occipital lobe, a target mask consisting of a coronal slice of the other occipital lobe, a coronal exclusion mask at approximately the level of the central sulcus, and a sagittal exclusion mask between the occipital lobes were used to reconstruct the forceps major (Fig. 1p). The forceps minor recipe was similar, with target and seed masks in coronal sections of the prefrontal cortex (Fig. 1q).

#### Anterior Commissure

A seed was placed at the midline of the anterior commissure and a target in the white matter between the globus pallidus and the putamen, based on descriptions from Dejerine and Dejerine-Klumpke [98]. An exclusion mask was placed superior to the seed and target masks and stop masks were placed posterior to the seed and target masks, to prevent streamline contamination from the rest of the basal ganglia, and in an axial section at the level of the extreme/external capsule, to prevent streamlines from going into the ventral pathway (Fig. 1r). Stop masks for the human (Fig. 2r) and macaque (Fig. 3r) were placed in slightly different orientations owing to the different spatial relationship of the basal ganglia and anterior commissure in these species.

#### Middle Cerebellar Peduncle

A seed was placed in the white matter stem of one cerebellar hemisphere and a target in the opposite (Fig. 1s). An exclusion mask was placed sagittally between the two cerebellar hemispheres and axially at the base of the thalamus. (Fig. 1s).

### Thalamic projections

#### Thalamic Radiations

Anterior thalamic radiation recipes consisted of an axial seed in the anterior third of the thalamus [99], with a target in the white matter of the prefrontal cortex just anterior to the basal ganglia, in the anterior thalamic peduncle (Fig. 1t). An exclusion mask was placed posterior to the thalamus. The superior thalamic radiation recipe consisted of a seed in the superior half of the thalamus, and a target in the pre- and postcentral gyri. Exclusion masks were placed coronally, posterior to the thalamus and anterior to the precentral gyrus (Fig. 11u). To reconstruct the posterior thalamic radiation, a seed in the posterior thalamus and a coronal mask in the occipital lobe was used (Fig. 1v). Exclusion masks were placed anterior and inferior to the thalamus (Fig. 1v).

#### Acoustic and Optic Radiations

For the acoustic and optic radiations, seeds were placed in the medial geniculate nucleus and lateral geniculate nucleus of the thalamus, respectively. A target was placed in the planum temporale for the acoustic radiation and in and around the calcarine sulcus for the optic radiation (Fig. 1w and 1x). Because of the different morphometry of macaque primary visual cortex, for this species the target mask was placed as a coronal section at approximately the level of the lunate sulcus (Fig. 3x).

### Surface projection maps

Group-level cortical surface representations of each tract were created to establish which cortical territories are reached. Since there are known issues of gyral bias and superficial white matter [14,68] when tracing towards the cortical grey matter, we employed a recent approach that aims to address some of these issues by multiplying the tractography results with a whole-brain connectivity matrix [8,20].

Tractography of the tracts was done as described above and the resulting tracts were warped to standard space, normalized, averaged, and thresholded. When combining several tracts, this effectively results in a (tract) x (brain) matrix. This tractogram matrix was then multiplied by a whole-brain (surface) x (brain) connectivity matrix derived from seeding in the mid-thickness surface and tracking to the rest of the brain. To keep computation time manageable, we generated this whole-brain connectivity matrix based on 10 individual surface reconstructions in the chimpanzee (32k vertices) and human (32k vertices), and for one common surface in the three macaques (10k vertices). To save computational load, the rest of the brain was downsampled (2 mm isotropic voxels for the human, 1.5 for the chimpanzee, 1 for the macaque).

To rebalance the weights in the tract surface projections to be more homogenous, values in the whole-brain connectivity matrix were weighted by the distance between vertex and voxel. A distance matrix across all vertices of the mid-thickness surface and all brain voxels was computed using MATLAB’s pdist2-function resulting in a matrix of the same size as the connectivity matrix. Each element in the whole-brain connectivity matrix was then divided by the corresponding value in the vertex-to-voxel distance matrix. To decrease data storage load (approximately 10 GB per whole-brain matrix) the weighted connectivity matrices of the subjects were averaged for each hemisphere and species. We then created the cortical surface representation by multiplying the (tract) x (brain) tractogram matrix with the weighted (surface) x (brain) connectivity matrix, resulting in a (tract) x (surface) matrix that we term the *connectivity blueprint [8]*. The columns of this blueprint represent the surface projection of each tract. Results were displayed onto one individual’s pial surface for chimpanzee and macaque, and onto an average surface in humans using Connectome Workbench [100]. Note that any relations to detailed individual surface landmarks should be interpreted with caution given that group-level results are displayed.

## Data availability

Analysis code, tractography recipes, and results will be made available online at locations linked from the lab’s website (www.neuroecologylab.org) upon acceptance of the paper. Analysis code is also part of the MR Comparative Anatomy Toolbox (Mr Cat; [101]). Raw data are available from the National Chimpanzee Brain Resource (www.chimpanzeebrain.org) for the chimpanzee, the PRIME-DE repository (http://fcon_1000.projects.nitrc.org/indi/indiPRIME.html) for the macaque, and from the Human Connectome Project (www.humanconnectome.org) for the human.

## Acknowledgements

K.L.B. was supported by a Marie Skłodowska-Curie Postdoctoral Research Fellowship from the European Commission [MSCA-IF 750026]; L. L. is supported by the National Institutes of Health [1R01EB027147, 2P50MH100029, 1R01MH118534, 1R01MH118285]; N.E. is a Wellcome Trust Doctoral student in Neuroscience at the University of Oxford [203730/Z/16/Z]. R.B.M. is supported by the Biotechnology and Biological Sciences Research Council UK [BB/N019814/1] and the Netherlands Organization for Scientific Research [452-13-015]. The Wellcome Centre for Integrative Neuroimaging is supported by core funding from the Wellcome Trust [203129/Z/16/Z]. Chimpanzee brain scans were acquired prior to the 2015 implementation of U. S. Fish and Wildlife Service and National Institutes of Health regulations governing research with chimpanzees, made possible through the Yerkes Base Grant [ORIP/OD P51OD011132]; an archive of these scans has been made available through the National Chimpanzee Brain Resource [NIH NINDS NS092988]. Human brain scans were made available through the Human Connectome Project [1U54MH091657].

## Notes

The authors would like to thank Shaun Warrington, Saad Jbabdi, and Stam Sotiropolous for their work on XTRACT toolbox and library (fsl.fmrib.ox.ac.uk/fsl/fslwiki/XTRACT) which helped enable this project. The authors would like to thank Guilherme Freches with his assistance on generating the chimpanzee whole-brain connectivity matrix.

---

**Figure S1.**
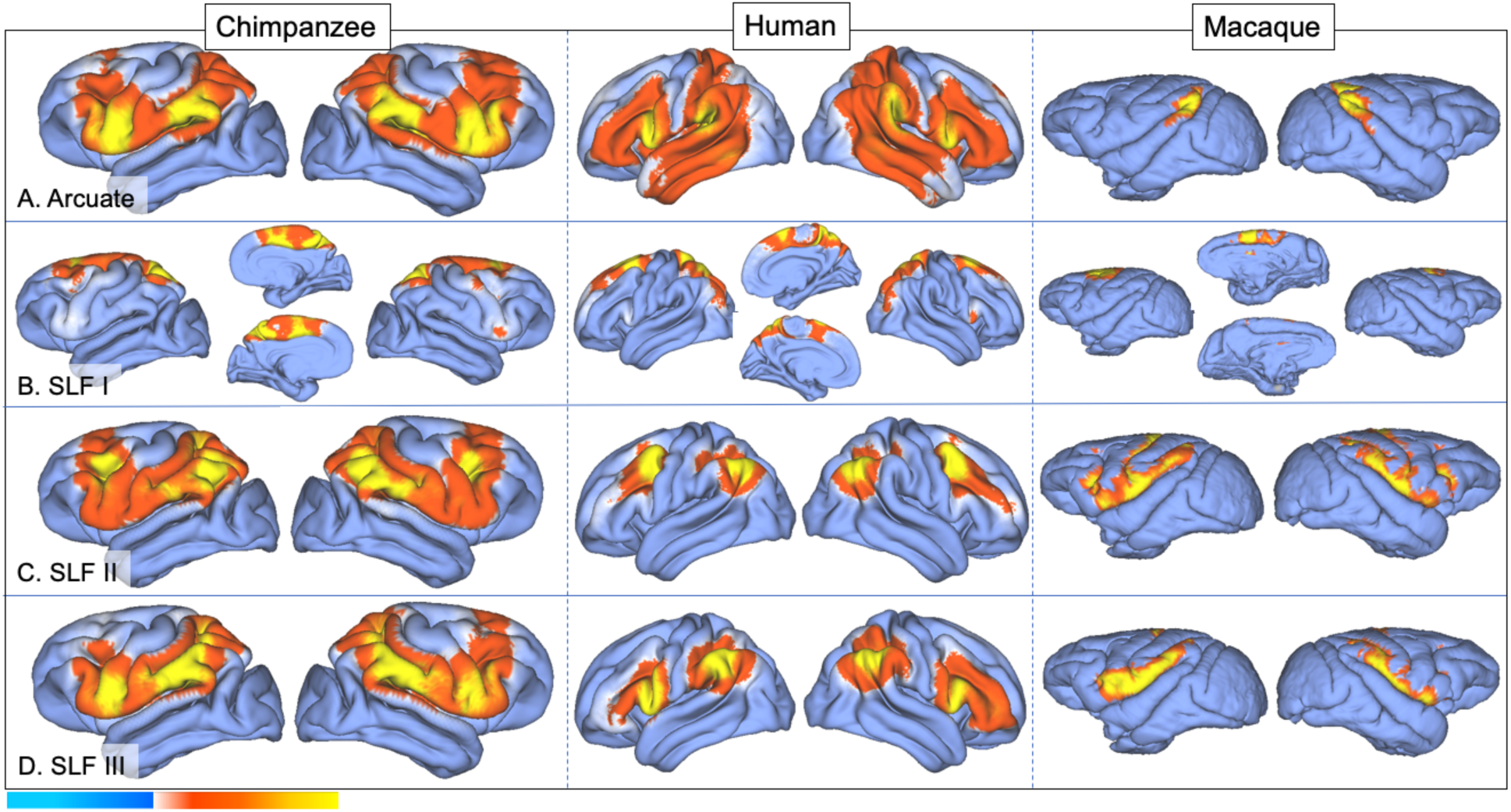
Surface projection results for arcuate (a) and SLFs I-III (b-d) in chimpanzee, human, and rhesus macaque. Color bar indicates heat map of tractogram normalized probability values.

**Figure S2.**
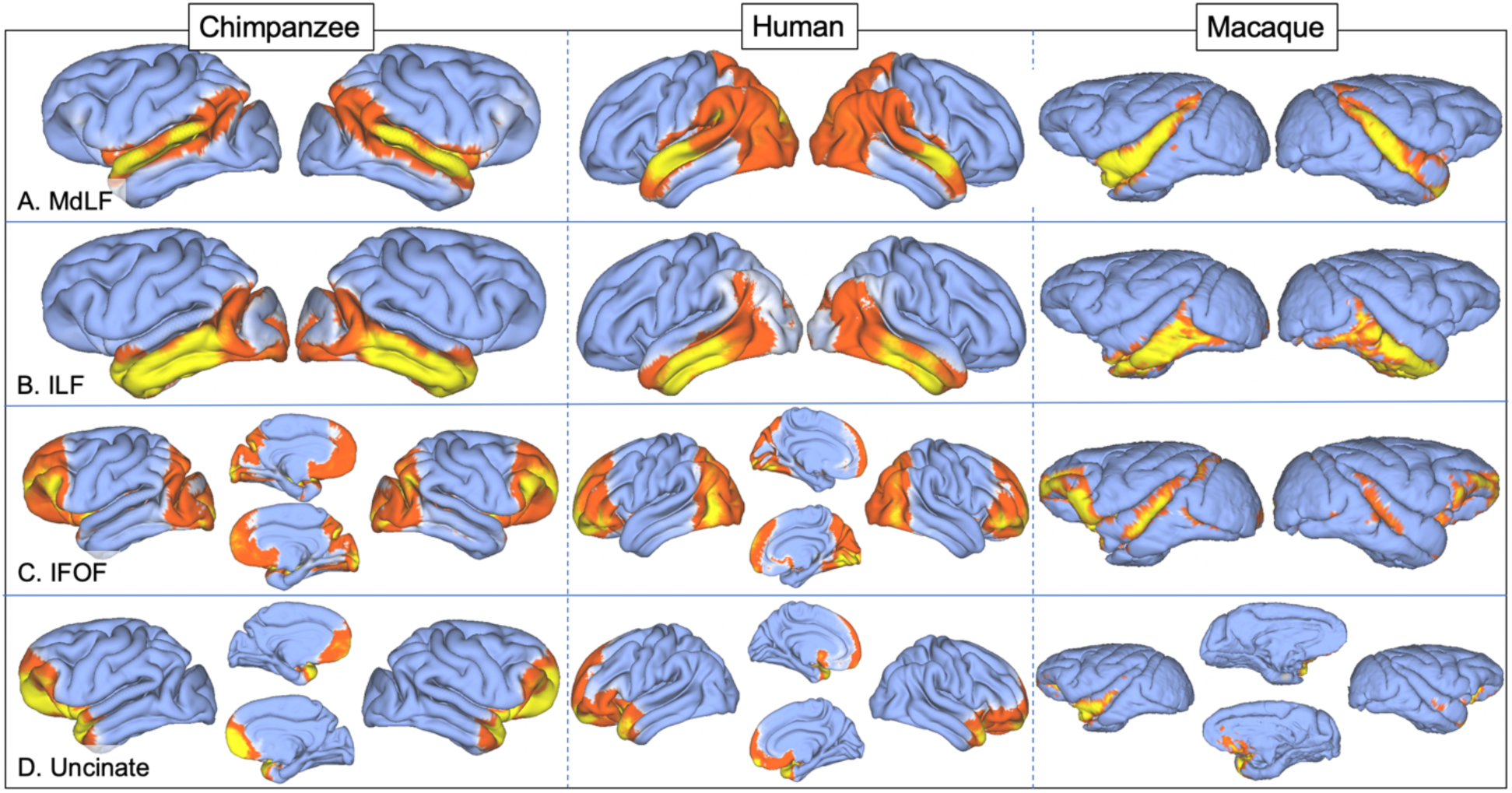
Surface projection results for MdLF (a), ILF (b), IFOF (c), and UF (d) in chimpanzee, human, and rhesus macaque.

**Figure S3.**
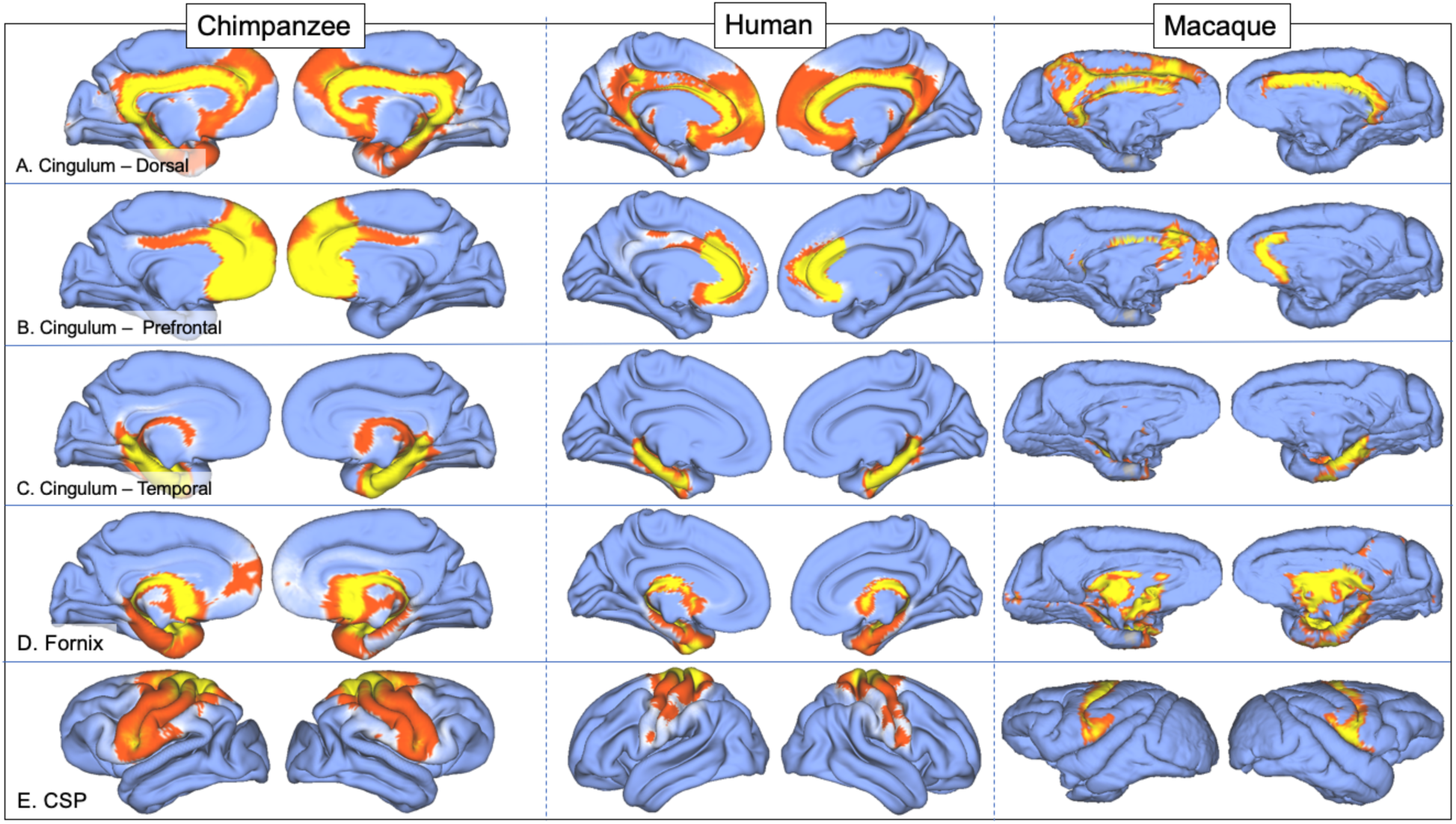
Surface projection results for cingulum bundle (a-c), fornix (d), and CST (e) in chimpanzee, human, and rhesus macaque.

**Figure S4.**
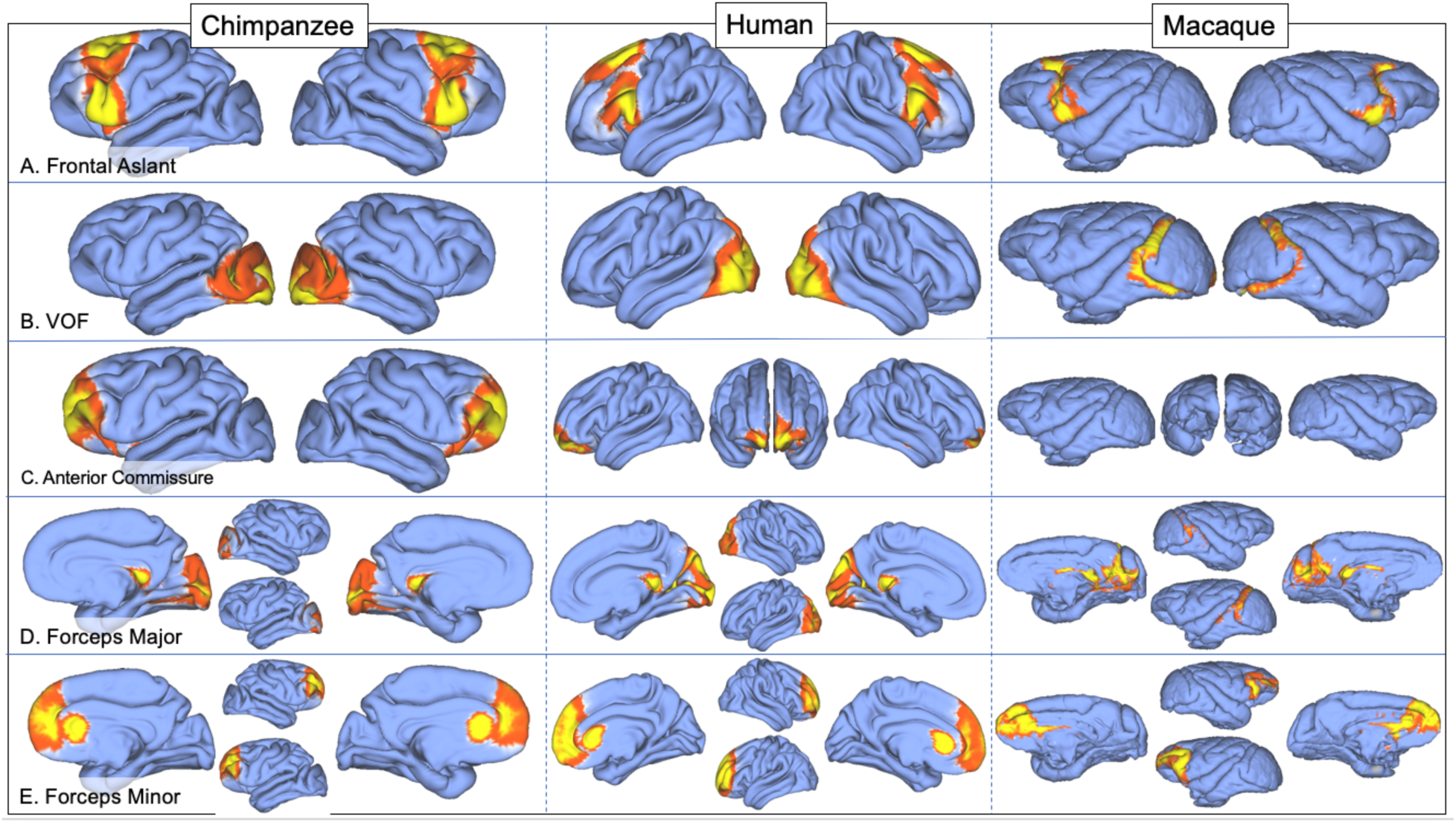
Surface projection results for frontal aslant (a), VOF (b), anterior commissure (c), and forceps major and minor (d, e) in chimpanzee, human, and rhesus macaque.

**Figure S5.**
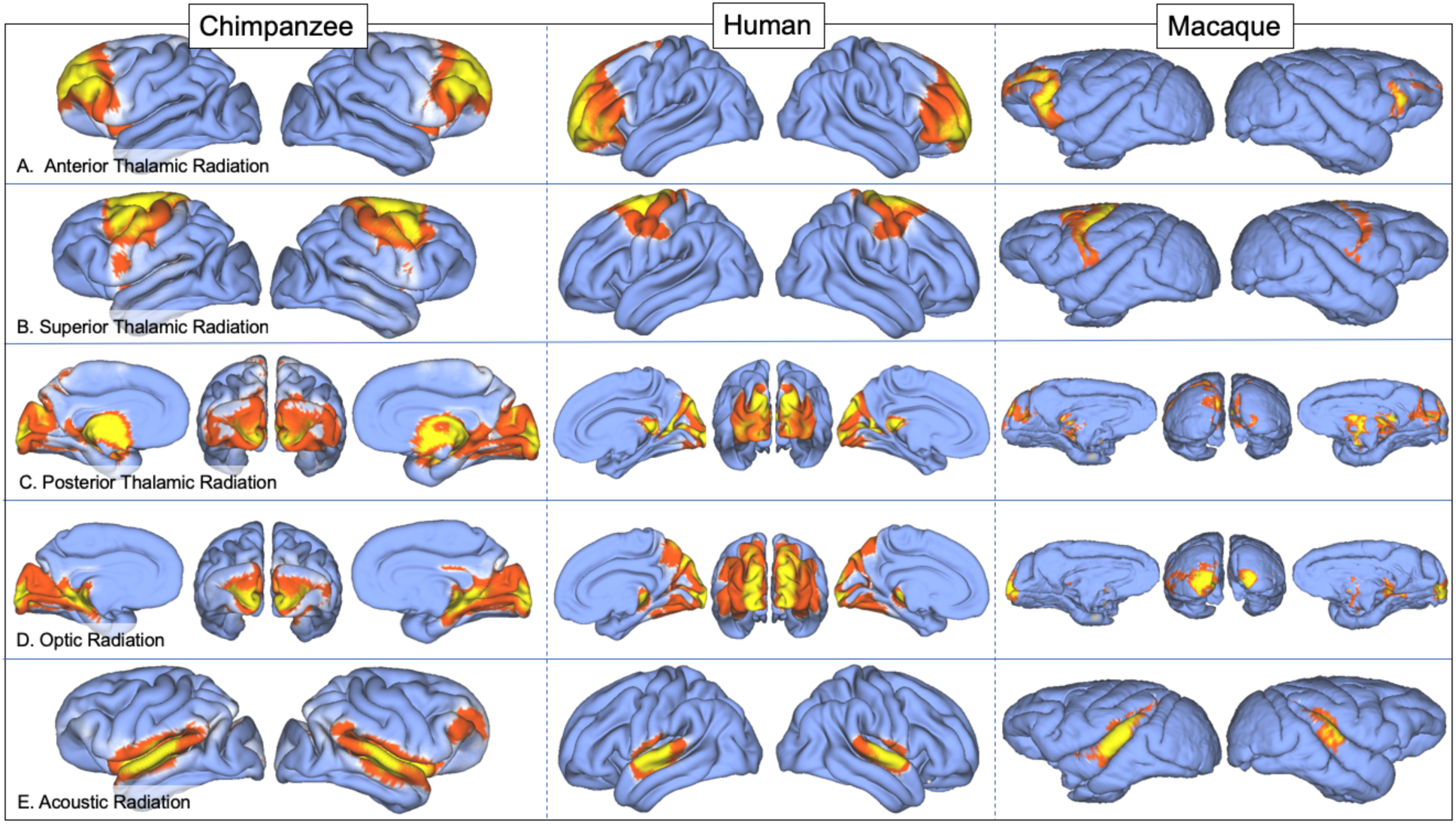
Surface projection results for anterior, superior, and posterior thalamic radiations (a-c), optic radiation (d), and acoustic radiation (e) in chimpanzee, human, and rhesus macaque.

**Figure S6:**
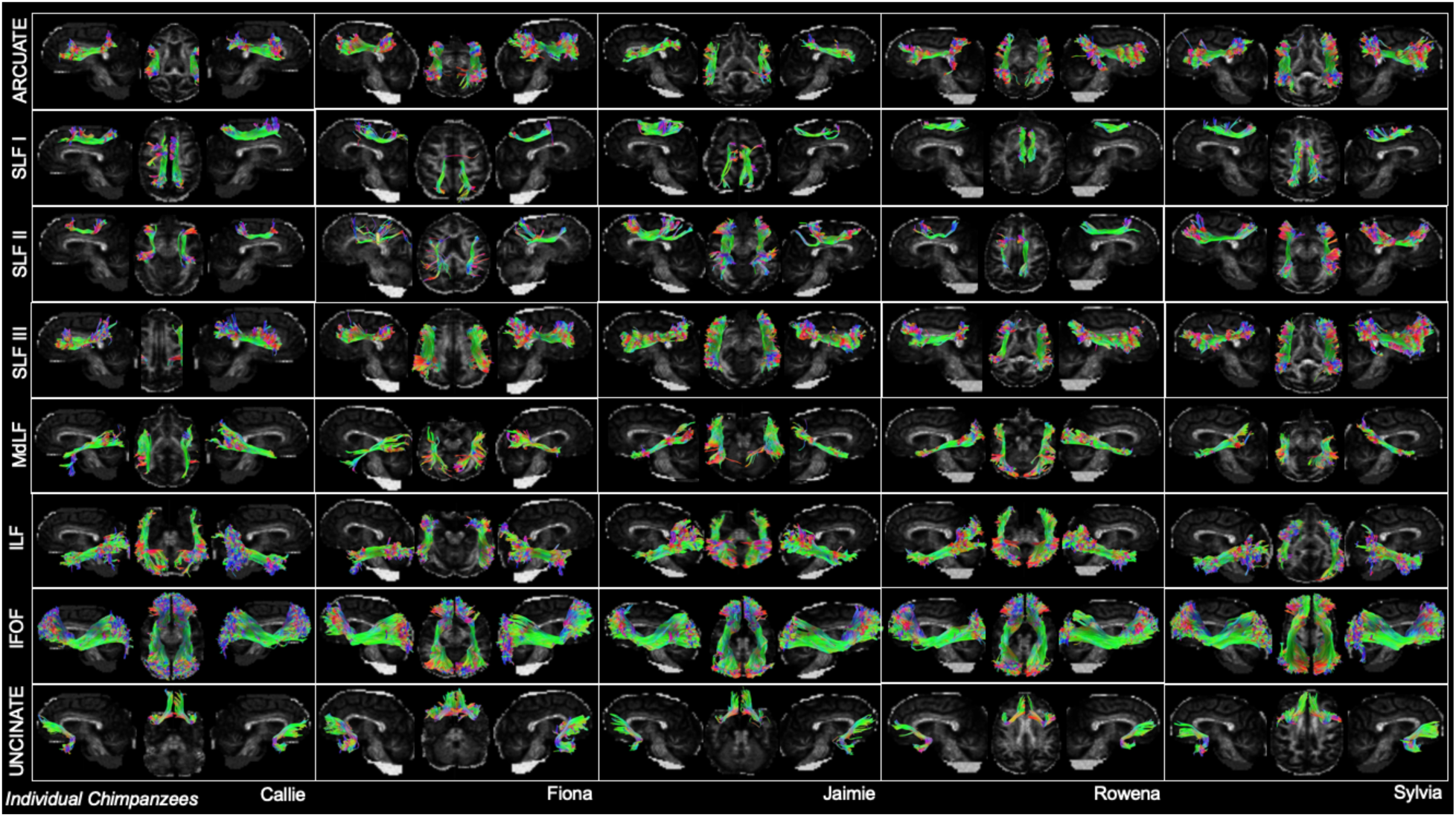
Deterministic tractography of large white matter tracts in a sampling of 5 chimpanzee individuals.

